# Detailed Regulatory Interaction Map of the Human Heart Facilitates Gene Discovery for Cardiovascular Disease

**DOI:** 10.1101/705715

**Authors:** Valerio Bianchi, Geert Geeven, Nathan Tucker, Catharina R.E. Hilvering, Amelia W. Hall, Carolina Roselli, Matthew C. Hill, James F. Martin, Kenneth B. Margulies, Patrick T. Ellinor, Wouter de Laat

**Author notes:** These authors contributed equally (Valerio Bianchi, Geert Geeven, Nathan Tucker).

## Abstract

Most disease-associated variants identified by population based genetic studies are non-coding, which compromises finding causative genes and mechanisms. Presumably they interact through looping with nearby genes to modulate transcription. Hi-C provides the most complete and unbiased method for genome-wide identification of potential regulatory interactions, but finding chromatin loops in Hi-C data remains difficult and tissue specific data are limited. We have generated Hi-C data from primary cardiac tissue and developed a method, peakHiC, for sensitive and quantitative loop calling to uncover the human heart regulatory interactome. We identify complex CTCF-dependent and -independent contact networks, with loops between coding and non-coding gene promoters, shared enhancers and repressive sites. Across the genome, enhancer interaction strength correlates with gene transcriptional output and loop dynamics follows CTCF, cohesin and H3K27Ac occupancy levels. Finally, we demonstrate that intersection of the human heart regulatory interactome with cardiovascular disease variants facilitates prioritizing disease-causative genes.

## Introduction

Genome-wide association studies (GWAS) have been remarkably successful at identifying the genetic basis of common human diseases and traits. However, the identification of target genes and underlying mechanisms remains challenging. Indeed, more than 90% of disease-associated variation resides in non-coding DNA (1000 Genomes Project Consortium et al., 2010; Chen et al., 2016; Maurano et al., 2012). Integration of chromatin features has demonstrated that GWAS risk variants are enriched in evolutionary conserved elements, regions of open chromatin, and potential tissue-specific regulatory elements (Maurano et al., 2012; Roadmap Epigenomics Consortium et al., 2015; Schaub et al., 2012). These findings suggest that GWAS-associated variants may affect disease risk by changing the function of distal regulatory elements controlling gene expression.

To act on gene promoters, regulatory elements need to be in close spatial proximity. Distant regulatory elements can accomplish this through chromatin looping (Deng et al., 2012; Tolhuis et al., 2002). This articulates the ‘3D genome promise’, which hypothesizes that the identification of all chromatin loops in all cell types will link distal regulatory elements to genes and thus help assign function to non-coding genetic variation in health and disease (Krijger and de Laat, 2016; Spielmann et al., 2018). In recent years, various studies have emphasized the relevance of utilizing genome topology for the interpretation of disease-associated variation (Gröschel et al., 2014; Liu et al., 2017; Lupiáñez et al., 2015; Smemo et al., 2014). For example, a study that employed 4C-seq, a chromosome conformation capture technique, shifted the paradigm for the genetic locus most significantly associated with obesity. The obesity-associated intronic variants were presumed to interact with the promoter of the nearest gene, *FTO*, but instead were found to interact over a long distance with the homeobox gene *IRX3* (Smemo et al., 2014). In subsequent work, the obesity risk variant was shown to disrupt a repressor binding site, which led to enhancer release and doubling of *IRX3,* but not *FTO* expression, as well as increased lipid storage in early adipocyte differentiation (Claussnitzer et al., 2015).

Chromosome conformation capture (3C) methods (Dekker et al., 2002), and in particular, the high throughput variants 4C, 5C, and Hi-C (Dostie et al., 2006; Lieberman-Aiden et al., 2009; Rao et al., 2014; Simonis et al., 2006), have proven instrumental for uncovering genome structures inside the living cell nucleus. They demonstrated that long-range contacts between enhancers and promoters are often cell-type specific and typically confined within topologically associating domains (TADs). TADs are looped structures inside chromosomes that span an average size of 1 Mb and that contribute to cell-type specific gene activity, replication timing and chromatin signatures (Bonev et al., 2017; Dixon et al., 2012; Nora et al., 2012). The regulatory function of TADs has become clear in recent studies, where disruption of TAD boundary loops was observed to cause deregulation of genes that can lead to disease (Krijger and de Laat, 2016; Nora et al., 2012; Rao et al., 2017; Spielmann et al., 2018). Thus, to systematically interrogate the topological and possible functional consequences of non-coding genetic variation, it is necessary to assess their involvement in all categories of chromatin loops, in the relevant cell type.

Promoter-enhancer interactions and sub-domain structures inside TADs are challenging to systematically detect from Hi-C data since they are more dynamic and tissue-specific than TAD-spanning loops. For this reason, targeted DNA conformation capture methods are currently preferred to intersect GWAS hits with chromatin loops. Examples of such methods are promoter-capture (Choy et al., 2018a; Hughes et al., 2014; Mifsud et al., 2015) and HiChIP strategies (Mumbach et al., 2017), which enrich Hi-C libraries for presumed relevant DNA interactions to enable their selective sequencing and analysis. These methods are inherently limited to the targeted regions of the genome and are more susceptible to technical bias, so they are less comprehensive and conclusive than Hi-C. Given these challenges, it would be ideal to develop methods for much more sensitive chromatin loop calling from Hi-C datasets.

Here we present peakHiC, an approach that extracts individual contact profiles for all genomic sites from replicate Hi-C datasets to search for reciprocally confirmed chromatin contacts and to quantify their interaction strength. peakHiC identifies an order of magnitude more loops than other approaches and finds genes, enhancers, and CTCF occupied sites in the genome that can be confirmed as valid and functionally relevant. We then generated Hi-C data from primary cardiac tissue and applied peakHiC to uncover a detailed spatial regulatory interactome of the human heart. Finally, we interpreted GWAS variants in the context of the folded genome, to greatly enhance the identification of disease associated genes at cardiovascular disease risk loci (Figure 1A).

**Figure 1.**
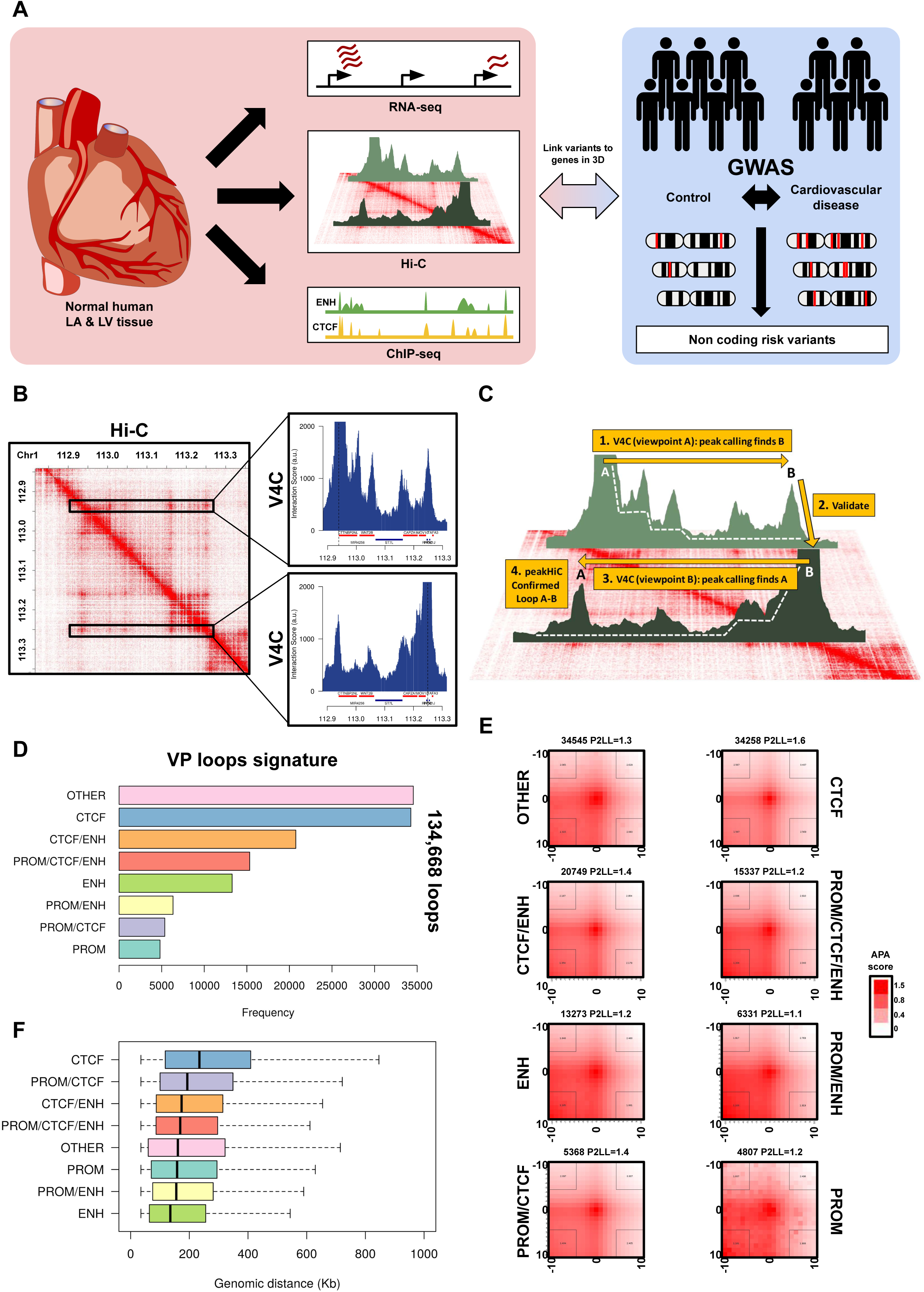
Chromatin loop identification by peakHiC. A) Experimental design. Heart tissue from human donors is used to generate transcriptome, epigenome and Hi-C 3D genome data to call and characterize cardiac chromatin loops and link non-coding cardiovascular disease-associated variants to genes and regulatory DNA elements. B) Heatmap of GM12878 Hi-C contacts (at 1Kb resolution), from which two virtual 4C contact plots are extracted for the gene promoters of CTTNBP2NL (top) and RHOC (bottom) that reciprocally contact each other. C) Principles of peakHiC. For a loop to be scored by peakHiC, both anchors must independently and reproducibly identify each other as a contact peak in their respective virtual 4C contact profile over its corresponding background contact profile. D) peakHiC called loops in GM12878, categorized by having their 10Kb viewpoint anchor overlapping with a gene promoter and/or a CTCF ChIP-seq peak and/or an H3K27Ac ChIP-seq peak. E) Aggregate Peak Analysis (APA) performed with Juicer on the categorized 134,629 GM12878 peakHiC loops sites. F) Boxplots showing the distribution distances spanned by each category of loops.

## Results

### Chromatin loop identification by peakHiC

To investigate the spatial configuration of the genome in primary human heart tissue, we applied *in situ* Hi-C to primary human left atrium (LA) and ventricle (LV) from four donors (**Table 1**). Using the frequent cutting restriction enzyme DpnII, an average of ∼385M valid Hi-C read pairs per sample were obtained, totaling 1.51B and 1.58B Hi-C contacts for LA and LV, respectively. Since the LA and LV Hi-C contact matrices were highly similar (see below), we combined all libraries to have an extensive human heart Hi-C dataset of >3B contact pairs. For comparison, we used the published deeply sequenced Hi-C dataset of the human lymphoblastoid cell line GM12878 (∼4.9B Hi-C contacts across 9 replicate Hi-C datasets: (Rao et al., 2014)).

Hi-C data can be transformed into so-called virtual 4C (V4C) contact profiles that visualize the chromatin interactions of given chromosomal sites of interest (‘viewpoints’, as generated by 4C technology (Van De Werken et al., 2012). For robustness, we extract, plot, and collectively analyze all contact pairs involving a bin (average size of ∼12 kb) of neighboring restriction fragments centered on a site of interest (gene promoter, enhancer, domain boundary, risk variant) (see methods and **Table 2** for further justification). Such V4C profiles may not show structural domains as clearly as traditional Hi-C contact matrices, but do provide a more intuitive and more detailed visualization of the contacts made by these genomic sites, at a resolution that we show is high enough to interpret their underlying principles (Figure 1B).

On initial visual inspection of overlays from our V4C contact profiles we found that many tissue-specific gene promoters had specific local chromatin interactions in the corresponding tissue (Figure S1A **and** S1B). Current Hi-C data analysis packages fail to systematically identify local but possibly functionally meaningful genomic interactions. We therefore developed an analytical tool to better enable mining Hi-C data for possibly regulatory chromatin loops.

We recently presented a non-parametric regression-based analysis tool, called peakC, for reproducible calling of chromatin interaction peaks from 4C and Capture-C datasets (Geeven et al., 2018). To create peakHiC, we modified this peak caller for systematic chromatin loop identification from our Hi-C derived V4C contact profiles (See Figure 1C). For chromatin loop identification by peakHiC, in principle any genomic site of interest can be interrogated. As viewpoints in this study, we selected the 49,052 currently annotated transcription start sites (TSSs) of coding and non-coding genes of the human genome (hg19) derived from the annotation database *TxDb* (UCSC) available on Bioconductor, complemented with a set of 493,937 putative CTCF binding sites derived from the CTCFBSDB 2.0 database (Ziebarth et al., 2013). Peak calling was limited to the 2Mb genomic interval surrounding each viewpoint, the interval in which the vast majority of regulatory sequences are located that have direct measurable impact on a gene’s expression (Bhatia and Kleinjan, 2014). For each viewpoint and replicate V4C profile an independent background model was calculated and used to rank contacts according to their signal over background. Rank-product statistics was applied across replicates to assign a P-value that evaluates the statistical significance of each identified loop. For a contact to be definitively scored as a loop anchor, we required that it had to similarly identify the original viewpoint (Figure 1C). We noticed that peakHiC-scored interactions largely corresponded to the visually appreciable peaks in V4C contact profiles (Figures S1A **and** S1B), which is the first requirement of a peak caller. peakHiC currently ignores interactions spanning less than 30kb, as loops within this range cannot reliably be discerned from the high local contact background.

To validate the method, we initially applied peakHiC to published GM12878 Hi-C datasets. peakHiC identified ∼135,000 high confidence chromatin loops in GM12878. This includes ∼83% of the loops called within 2Mb intervals by the widely used Hi-C analysis package HICCUPS (Durand et al., 2016), but gives a ∼15-fold increase in loop numbers (Figure S1C). To validate the newly identified chromatin loops, we applied aggregate peak analysis (APA), a critical quality check for contact specificity (Rao et al., 2014). The newly called loops showed clear focal contact enrichment in APA (Figure S1D), which confirmed their interaction specificity. Loop formation occurred over all distances with an efficiency that decayed exponentially (Figure S1E). Ninety-two percent of the peakHiC loops occurred inside previously called TADs (Dixon et al., 2012). We subdivided the loops into eight categories based on viewpoint characteristics, categorized based on either being promoter centered, CTCF-bound, H3K27Ac marked or combinations thereof, plus a category that had none of these characteristics at the viewpoint (Figure 1D). The majority of the loops were CTCF centered, but nearly 60,000 loops had no CTCF associated with the viewpoint. A large cohort of ∼35,000 loops had none of the aforementioned marks (Figure 1D). ChromHMM analysis applied to the GM12878 epigenome (Ernst and Kellis, 2012) showed that the most frequent signature on those anchors is quiescent or repressed chromatin (Figure S1F). All categories of loops showed clear focal contact enrichment in APA (Figure 1E), supporting they are valid loops. In general, CTCF-centered loops spanned the largest distances (mean 291Kb) while loops centered on enhancers acted over shorter genomic intervals (mean 191Kb) (Figure 1F). To further validate peakHiC loops, we next subjected the different categories to a more detailed investigation.

### The CTCF looping landscape uncovered by peakHiC in GM12878 cell line

We first focused on CTCF-anchored loops. PeakHiC found over 60% of all CTCF-bound sites involved in one or more chromatin loops in the GM12878 cell line (Figure 2A). Looped sites generally had higher CTCF- and cohesin ChIP-seq signals than non-looped CTCF occupied sites (Figure S2A). Cohesin-dependent CTCF looping is critical for domain formation but is also hypothesized to counteract the spatial aggregation of active and inactive TADs in A and B compartments (Haarhuis et al., 2017; Nuebler et al., 2018; Schwarzer et al., 2017) and prevent TAD compartment conversion (Schwarzer et al., 2017). While the overall CTCF and cohesin ChIP-seq signals were similar in A and B-type chromatin, both at looped and non-looped sites (Figure S2A), peakHiC found that a much larger percentage of all CTCF-bound sites in A than in B was engaged in looping (∼80% versus ∼50%) (Figure 2B). Since binding site density was also more than two-fold higher in A than B (i.e. in A more binding sites are accessible) (Figure S2B), CTCF looping density was more than 4-fold higher in the A than in the B compartment (∼54 versus ∼12 loops per Mb, in A and B, respectively) (Figure 2C). When considering all loops, we found a looping density (across the cell population) of ∼100 loops/Mb in the A compartment versus ∼20 loops/Mb in the B compartment (Figure 2C). This would agree with the idea that active loop formation is required to counteract heterochromatinization of active chromatin regions.

**Figure 2.**
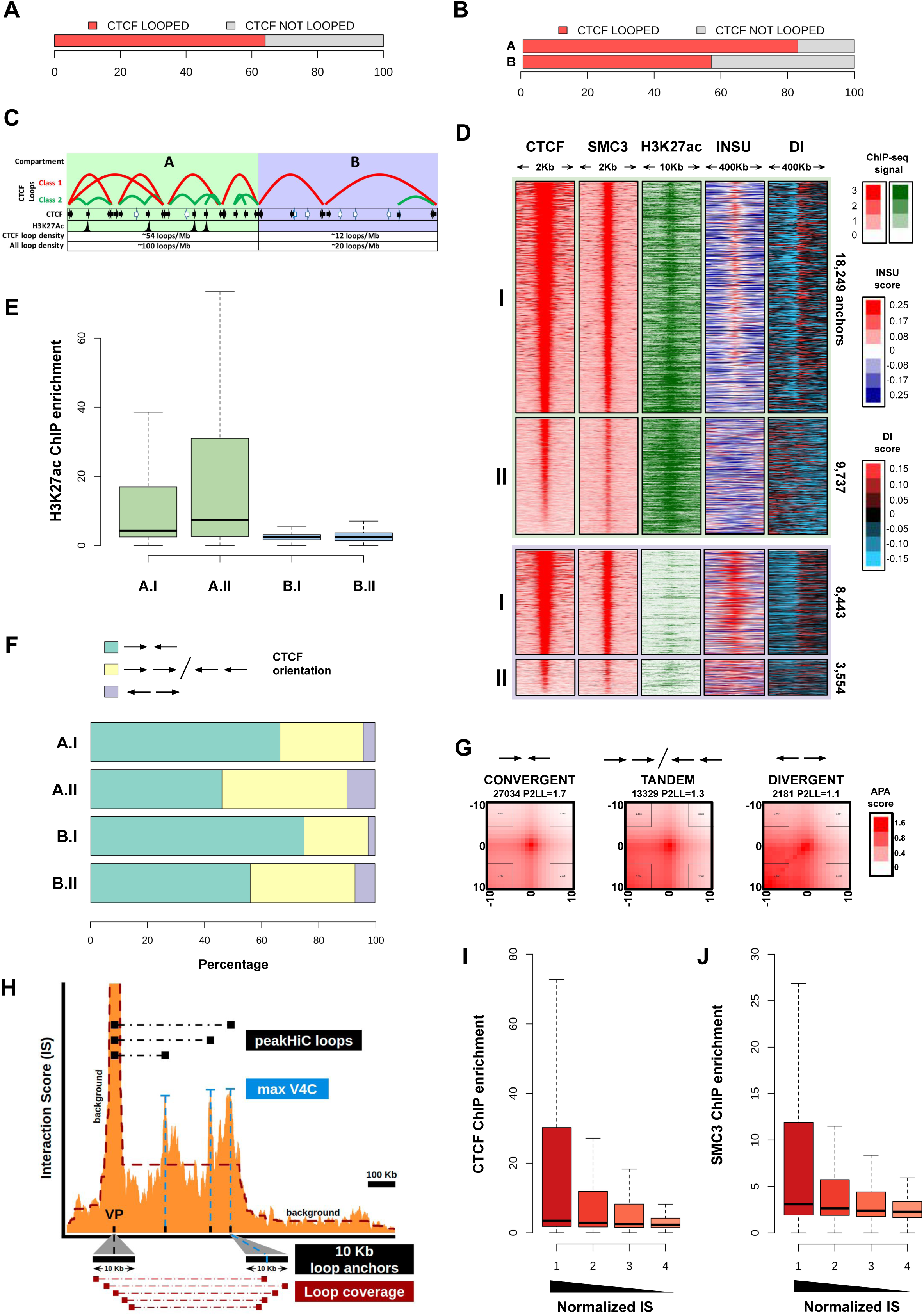
The CTCF looping landscape in GM12878. A) Barplot showing the percentage of all CTCF bound sites (as determined by ChIP-Seq) found engaged by peakHiC in chromatin looping, in GM12878. B) Barplots as in panel A, but now split based on whether the CTCF site is located in the “A” or “B” compartment. C) Schematic showing the density of class I and class II CTCF loops per megabase, in the A and the B compartment. D) Heatmaps of CTCF-bound chromatin loop anchors in the A (top) and B (bottom) compartment, ranked according to anchor-associated CTCF ChIP-seq enrichment, showing the SMC3 (cohesin) and H3K27ac ChIP-seq signal and the Hi-C derived insulation score and directionality index (DI) in GM12878 cells. Classification into type I and II loops is based on anchors showing (class I) or not showing (class II) a switch in DI (see Methods). E) Boxplots showing H3K27ac ChIP-seq enrichment at the anchors of class I and II loops, in the A and B compartment. F) Stacked barplot showing the relative contribution of pairs of convergent (green), tandem (yellow) and divergent (purple) CTCF bound sites to class I and class II loops, in the A and B compartment. G) Heatmap plots that show the result of aggregate peak analysis (APA), which measures normalized enrichment of Hi-C contacts between pairs of convergent (left), tandem (middle) and divergent (right) oriented CTCF-associated binding sites in GM12878 cells. All categories show enrichment, confirming the validity of these chromatin loops. H) Schematic showing how interaction scores are assigned to pairs of anchors forming a chromatin loop identified by peakHiC. Virtual 4C (V4C) contact profiles serve to link a viewpoint anchor to its contacting loop anchors, as described. The selected viewpoint site (e.g. a transcription start site (TSS) or a CTCF binding site) defines the center of the 10kb viewpoint bin, the summit of its contact peak (i.e. where its V4C score is maximal) defines the center of the 10kb contacting loop anchor bin. The sum of all contact events measured between any pair of restriction fragments located in these two 10kb bins, normalized for the complexity of the Hi-C library, defines the interaction score between the two loop anchors. I-J) Boxplots showing that CTCF (I) and SMC3 (J) protein occupancy levels at DNA loop anchors (based on ChIP-seq enrichment) in GM12878 correlate with the strength of their most prominent loop (as determined based on Hi-C interaction scores normalized for linear distance (see methods)).

In each compartment, we ranked the looped CTCF sites according to their CTCF ChIP-seq signal strength (Figure 2D). This ranking uncovered two distinct classes of CTCF-centered loops (see methods). Class I loops (∼65%) are centered on the sites that most efficiently recruit CTCF and cohesin. They display the typical hallmarks of domain boundaries: they insulate chromatin (i.e. block DNA interactions between flanking sequences) and mark sites where DNA contacts switch direction from preferentially upstream to downstream engagements (Figure 2D). Class I loops can therefore be classified as domain loops. Class II represents loops anchored on the ∼35% sites that less efficiently recruit CTCF and cohesin, display no obvious insulation capacity, and show no clear focal switch in contact directionality (Figure 2D). This may imply that insulation is not coupled to looping, or that class II loops occur in too few cells simultaneously to appreciate these associated features in cell populations. Class II CTCF loops are predominantly found in the A compartment (17326 of all 22704 class II loops), where they often have high levels of H3K27Ac at their anchors (Figure 2D **and** 2E). They span smaller distances than class I loops (median distance of 206Kb for class II versus 245 Kb for class I in the A compartment and 283 Kb versus 342 Kb in the B compartment, Figure S2C) and frequently (78%) share an anchor with a class I loop. Class II loops are therefore referred to as intra-domain loops. Over 50% of the class II loops represent interactions between non-convergent CTCF binding sites (Figure 2F), and APA confirmed these are loops with significant enrichment (APA P2LL between 1.1 and 1.7, Figure 2G).

A major advantage of peakHiC is that once chromatin loops have been identified, scores can be assigned to their interaction strength, allowing for a more quantitative analysis of Hi-C derived chromatin contact frequencies. We assigned an interaction score (IS) to each chromatin loop based on the number of observed Hi-C ligation events between the 10Kb region of the V4C viewpoint and the 10Kb region under the summit of the interaction peak (maximum V4C score) (see methods) (Figure 2H). To compare loop strengths across length scales, we converted this into distance-normalized interaction scores (see methods) and compared these quantitatively to CTCF- and cohesin-ChIP-seq signals. Assuming that ChIP-seq and Hi-C measures reflect the percentage of cells having these features, we hypothesized that ChIP-seq derived binding efficiency scores for CTCF and cohesin and Hi-C derived looping interaction scores should be correlated. Indeed, we observed a clear trend between Hi-C loop strength and median levels of ChIP enrichment for both CTCF and the cohesin subunit SMC3 (Figures 2I **and** 2J).

### Transcription regulatory interactions uncovered by peakHiC in GM12878 cell line

We then used peakHiC to identify loops centered on 22,918 promoters of coding genes. We found nearly 70% (15,600) of all gene promoters involved in a total of 48,705 chromatin loops (average ∼3 loops/promoter). 90% of promoter-centered loops formed within a TAD (intra-TAD loops) and collectively contacted ∼72% (31,293) of all H3K27Ac marked enhancers in GM12878 cells (Figure 3A). Most of the promoter containing loops (∼82%) had CTCF bound in at least one anchor, but 8,568 (∼18%) loops did not have CTCF bound at any of its anchors. APA analysis confirmed both categories represent true loops (Figure 3B), which highlights the existence of CTCF-independent loops.

**Figure 3.**
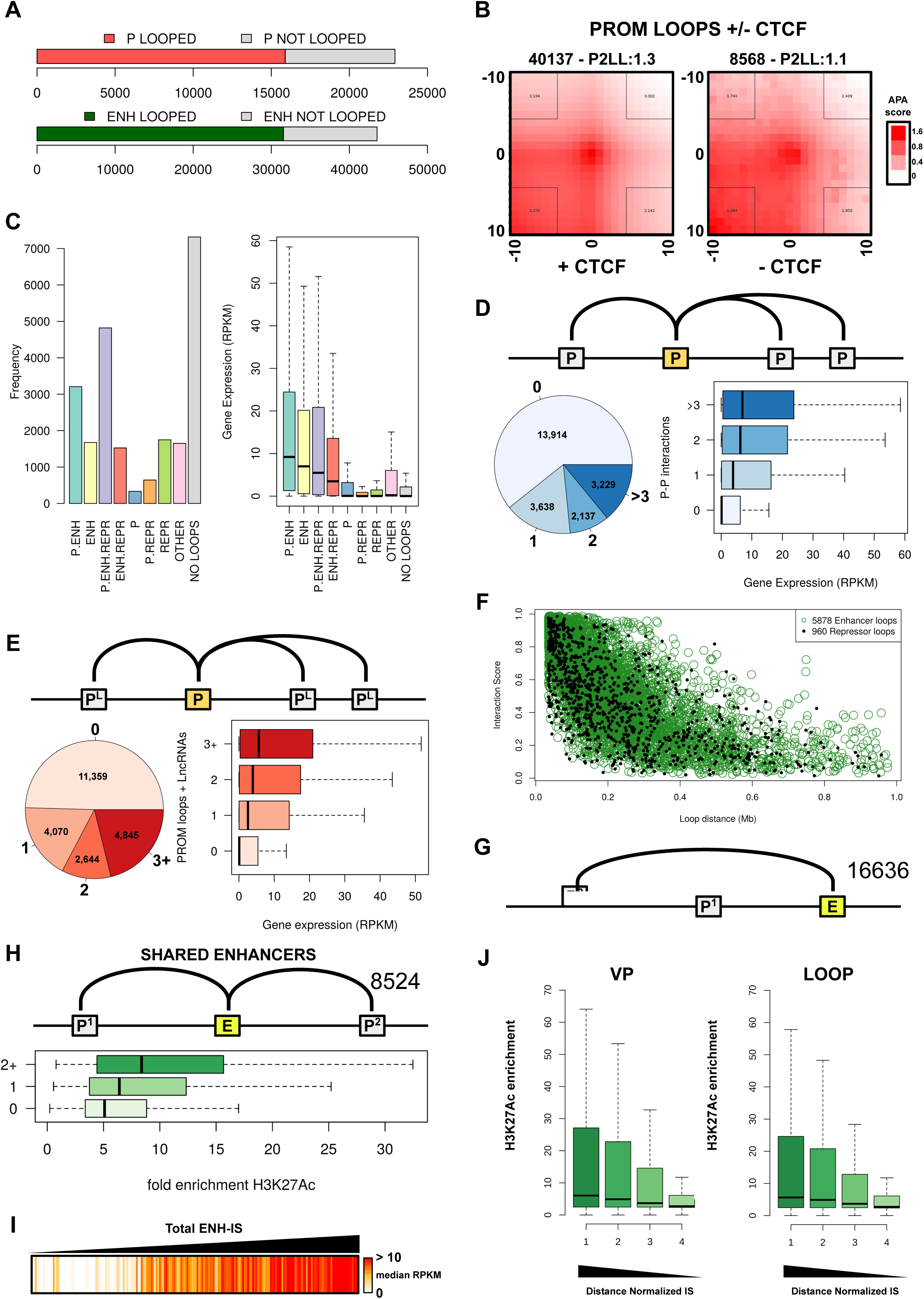
Transcription regulatory interactions uncovered by peakHiC in GM12878. A) Stacked barplot showing the proportion of gene promoters (top) and H3K27Ac-marked enhancers (bottom) involved in chromatin looping, as defined by peakHiC in GM12878 cells. B) Aggregate Peak Analysis (APA) applied to promoter-centered peakHiC loops with (left) or without (right) CTCF bound to any of its anchors. C) Left: barplot showing the not looped genes (NO LOOPS) and looped genes (from panel A), with the latter further categorized based on whether or not they contact another promoter (P) and/or having H3K27Ac ChIP-seq peaks (ENH) and/or a ChromHMM repressive chromatin signature at any of its loop anchors. Right: boxplot summarizing the gene expression (in RPKM) of genes belonging to each of these categories. D) Promoter-promoter contact networks. Pie-chart shows the distribution of promoters involved in contacts with different numbers of other promoters. Boxplot showing the gene expression (in RPKM) distribution for each of these categories of genes. E) Promoter-LncRNA promoter contact networks. Pie-chart shows the distribution of promoters involved in contacts with different numbers of LncRNA promoters. Boxplot showing the gene expression (in RPKM) distribution for each of these categories of genes. F) Interaction score (normalized) of promoter-centered enhancer (green) and repressor (black) loops, plotted against the linear distance spanned by the loops. Enhancer loops have strong H3K27Ac signal (ChIP-seq signal > 50th percentile) and low H3K27me3 signal (ChIP-seq signal < 1st percentile) at the anchors. Repressor loops have strong H3K27me3 signal (ChIP-seq signal > 50th percentile) and low H3K27Ac signal (ChIP-seq signal < 1st percentile) at the anchors. G) Illustration showing the number of promoter-enhancer loops (16323) that link an enhancer to a non-nearest promoter. H) Illustration shows the number of loops linking distinct promoters to shared enhancers. Boxplot shows the H3K27Ac ChIP-seq signal (log2 fold enrichment) at enhancers contacting two or more promoters (top), one promoter (middle) or no promoter (bottom): stronger enhancers more likely engage with multiple genes. I) Genome-wide quantitative correlation between the enhancer interaction score of genes and their transcriptional output. Genes are ranked from left to right according to increased total enhancer interaction score and are color-coded according to their transcription output in GM12878 cells (white = low, red = high expression). J) Stronger enhancers form stronger loops. Boxplots show the H3K27Ac ChIPseq signal distribution at anchors of loops grouped (per quartile) according to loop strength (distance normalized), at the viewpoint (left) and the contacting anchor (right).

A further inspection of the promoter-centered loops revealed a number of interesting loop categories. For example, we found many loops between promoters (P-P loops): roughly 60% of the looped gene promoters contacted at least one other gene promoter (Figure 3C **and** 3D). In fact, many complex P-P interaction networks were found involving three, four, or sometimes even more promoters. As was also suggested from ChIA-PET experiments (Li et al., 2012a), the degree of P-P loop connectivity correlated (Spearman’s rank correlation 𝛠=0.318) to the transcriptional output of the participating genes (Figure 3D). We additionally found a large collection of coding gene promoters (11,559) looping to long non-coding RNA (lncRNA) promoters (see below). Here the same trend was visible (Spearman’s correlation rhos=0.309): the more individual contacts a gene made with different lncRNA promoters, the higher its overall transcriptional output (Figure 3E). This supports the idea that promoters of coding and non-coding genes can also act as long-range enhancers of transcription⍰⍰.

An unexpected finding was that more than 50% of all promoters also loop to sites with a repressive epigenetic signature, as judged from the GM12878-derived ChromHMM profiles. Figure 3C classifies the genes according to their loop types in GM12878 cells (with enhancers (ENH), promoters (P), repressors (REPR), or combinations hereof) and shows that the transcriptional output was highest for genes contacting enhancers. Genes contacting repressors were generally inactive, as were genes not engaged in looping. Both enhancer and repressor loop occurred over all distance ranges (Figure 3F). Little is known about long-range gene repression. While our data may suggest that repression through looping is a common transcription regulatory mechanism, targeted disruption of these loops would be required for further investigation.

We focused on the promoter to enhancer (P-ENH) loops and quantified how frequently enhancers contacted the non-nearest gene. We found >16,600 enhancers doing this, which represents 98% of all enhancers engaged in promoter looping (Figure 3G). Whether these enhancers skipped the nearest gene or also contacted it was often unclear, as our method is unable to score loops within a 30 kb distance. Nevertheless, the data argue that nearest gene approaches are insufficient to comprehensively identify target genes of enhancers. Cellular genetic screens recently identified enhancers that are functionally acting on multiple genes simultaneously (Gasperini et al., 2019). Interestingly, we identified >8,500 loops connecting two or more gene promoters to shared enhancers. Enhancer strength (assessed from H3K27Ac fold enrichment) seemed to influence the ability to interact with more than one promoter, as the most complexly connected enhancers also had the highest H3K27Ac levels (Figure 3H).

Our ability to assign an interaction score to each chromatin loop enabled us to compute a total enhancer interaction score (IS) per gene, which we calculated by summing the interaction scores with individually contacted enhancers (totENH-IS, see methods). When we ranked all gene promoters according to their totENH-IS and intersected with the gene expression data, a striking genome-wide correlation between enhancer interaction frequencies and transcriptional output was found (Figure 3I). This strongly supports the idea that transcriptional activity is regulated through promoter-enhancer contact frequencies. Moreover, as seen before for CTCF loops, we observed an overall quantitative relationship between enhancer strength (i.e. H3K27Ac levels measured by ChIP-seq) and loop strength as measured by Hi-C in cell populations (Figure 3J).

In summary, we demonstrate a genome-wide relationship between enhancer contact frequencies and gene activity, and illustrate that loop strength correlates with enhancer strength. Additionally, peakHiC uncovers extensive promoter-promoter contact networks, functionally unrelated promoters that contact shared enhancers, promoters that contact repressive sites and promoters contacting lncRNA promoters. Based on interaction scores these loops can now be prioritized and systematically analyzed for their functional relevance.

### The architectural and regulatory chromatin loops of the human heart

Having validated the relevance of loops and interaction scores called by peakHiC on published Hi-C data from GM12878 cells, we applied the method to our Hi-C datasets generated from the left atrium (LA) and left ventricle (LV) of primary human heart tissue. For data interpretation, we also generated H3K27Ac ChIP-seq and RNA-seq profiles from the same donor tissue. We initially merged the LA and LV peakHiC loops, because we observed a very high correlation between LA and LV in Hi-C contacts at loops called in both conditions, even higher than between individual GM12878 datasets (Figure 4A **and** Figure S4A). We thus obtained a total of >3B of valid Hi-C read pairs (∼1.5B per heart tissue), which allowed calling a set of ∼130,000 heart loops by peakHiC.

Overall, very similar chromatin contact networks were identified in primary human heart tissue as in the GM12878 cell line. We defined heart A and B chromatin compartments based on clustering patterns in the heart Hi-C datasets (see methods). Class I and class II CTCF loops were readily appreciable based on the ranking of their CTCF ChIP-seq score (Figure 4B), with the class I domain loops having high CTCF occupancy and showing strong insulation at the anchors. Class II, the intra-domain loops, had low CTCF occupancy and showed no appreciable insulation at the anchors, but had high H3K27Ac levels specifically in the A compartment. In heart, ∼65% of gene promoters (14,940) were found involved in looping, collectively contacting ∼66% of the heart enhancers (27,535) (Figure 4C). We used our H3K27Ac ChIP-seq data in combination with ChromHMM data from heart to classify the looped gene promoters. Also, in heart the overall transcriptional output was highest for genes looped to enhancers, while gene promoters exclusively looped to repressive chromatin sites had uniform low expression (Figure 4D). Furthermore, as in GM12878 cells, we found a genome-wide correlation between the transcriptional output of individual genes and their total enhancer interaction score (Figure 4E), and between enhancer loop strength and H3K27Ac accumulation at loop anchors (Figure 4F). A significant number (10,224) of genes looped to lncRNA promoters, with some genes showing high expression in relation to high contact frequencies but without this being a genome-wide trend (Figure 4G). Nearly 8,000 promoters were spatially connected to at least one other gene promoter and again the degree of connectivity correlated with the expression levels of the participating genes (Figure 4H). Collectively, our findings from human heart tissue support our observations from GM12878 data and demonstrate that similar regulatory contact networks are appreciable by Hi-C applied to proliferating tissue culture cells and frozen primary human tissues.

**Figure 4.**
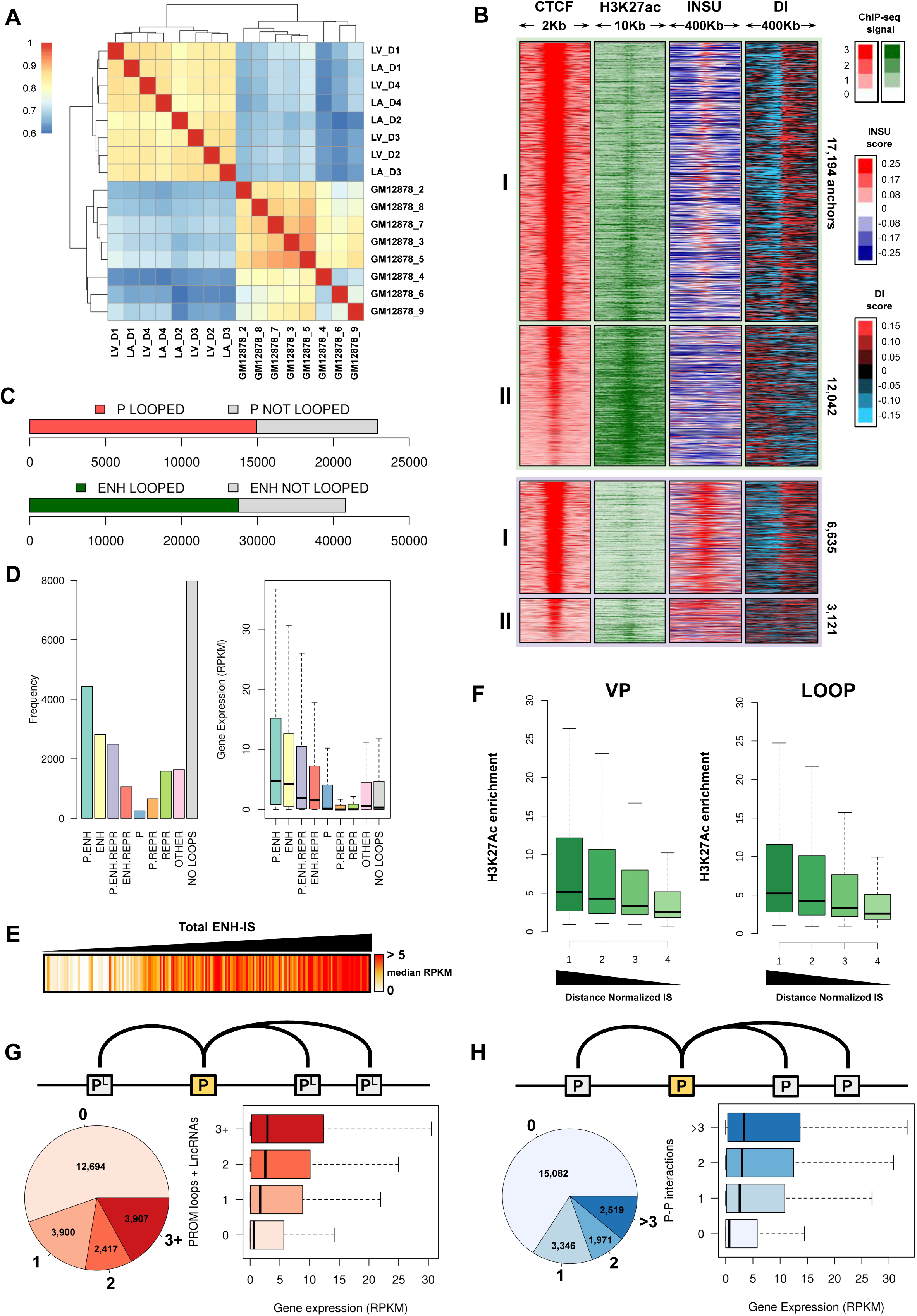
The architectural and regulatory chromatin loops of the human heart. A) Similarities between peakHiC loop strength measured in human left atrium (LA) and left ventricle (LV) heart tissue and GM12878 cells. Spearman’s rank correlation was computed for all independent Hi-C templates in LV (4 replicates), LA (4 reps) and GM12878 (8 reps). B) Heatmaps of CTCF-bound chromatin loop anchors in the A (top) and B (bottom) compartment, ranked according to anchor-associated CTCF ChIP-seq enrichment, showing the anchor-associated H3K27ac ChIP-seq signal, Hi-C derived insulation score and directionality index (DI), as measured in human heart tissue. Classification into type I and II loops is based on anchors showing (class I) or not showing (class II) a switch in DI (see Methods). C) Stacked barplot showing the proportion of gene promoters (top) and H3K27Ac-marked enhancers (bottom) involved in chromatin looping, as defined by peakHiC in human heart. D) Left: barplot showing the not looped genes (NO LOOPS) and looped genes categorized based on whether or not they contact another promoter (P) and/or having H3K27Ac ChIP-seq peaks (ENH) and/or a ChromHMM repressive chromatin signature at any of its loop anchors in heart. Right: boxplot summarizing the gene expression (in RPKM) of genes belonging to each of these categories. E) Genome-wide quantitative correlation between the enhancer interaction score of genes and their transcriptional output. Genes are ranked from left to right according to increased total enhancer interaction score and are color-coded according to their transcription output in heart tissues (white = low, red = high expression). F) Stronger enhancers form stronger loops. Boxplots show the H3K27Ac ChIPseq signal distribution at anchors of loops grouped (per quartile) according to loop strength (distance normalized), at the viewpoint (left) and the contacting anchor (right). G) Promoter-LncRNA promoter contact networks. Pie-chart shows the distribution of promoters involved in contacts with different numbers of LncRNA promoters. Boxplot shows the gene expression (in RPKM) distribution for each of these categories of genes in heart. H) Promoter-promoter contact networks. Pie-chart shows the distribution of promoters involved in contacts with different numbers of other promoters. Boxplot shows the gene expression (in RPKM) distribution for each of these categories of genes in heart.

### Tissue specific chromatin loops of the human heart

We then sought to identify and characterize the most heart- and GM12878-specific chromatin loops. For this, we first merged all heart and GM12878 datasets to obtain a total set of 234,838 loops (see methods). We classified them, based on the characteristics of their viewpoints in heart, into the eight previously described categories. We first investigated loop dynamics by simply interrogating whether each loop was called in GM12878, heart, or in both cell types. Loops with CTCF bound at their viewpoint anchor were more conserved between the tissues (44% conserved, 44,398) than loops not having CTCF at their viewpoint (26% conserved, 34,016 loops) (Figure 5A **and** Figure S5A). Class I domain loops were overrepresented among the tissue-invariant CTCF loops, while class II intra-domain CTCF loops were overrepresented among the tissue-specific CTCF loops (Figure 5B). To more quantitatively assess loop dynamics, we ranked the loops according to the difference in interaction scores between heart tissue and GM12878 cells. All categories of loops proportionally contributed to the 5% collection of most heart-specific loops (Figure 5C). We re-classified the loops based on GM12878-specific features and repeated the analysis to identify the 5% most GM12878-specific chromatin loops. Among these, enhancer-centered loops were over-represented and CTCF-anchored loops were underrepresented (Figure S5A **and** S5B).

**Figure 5.**
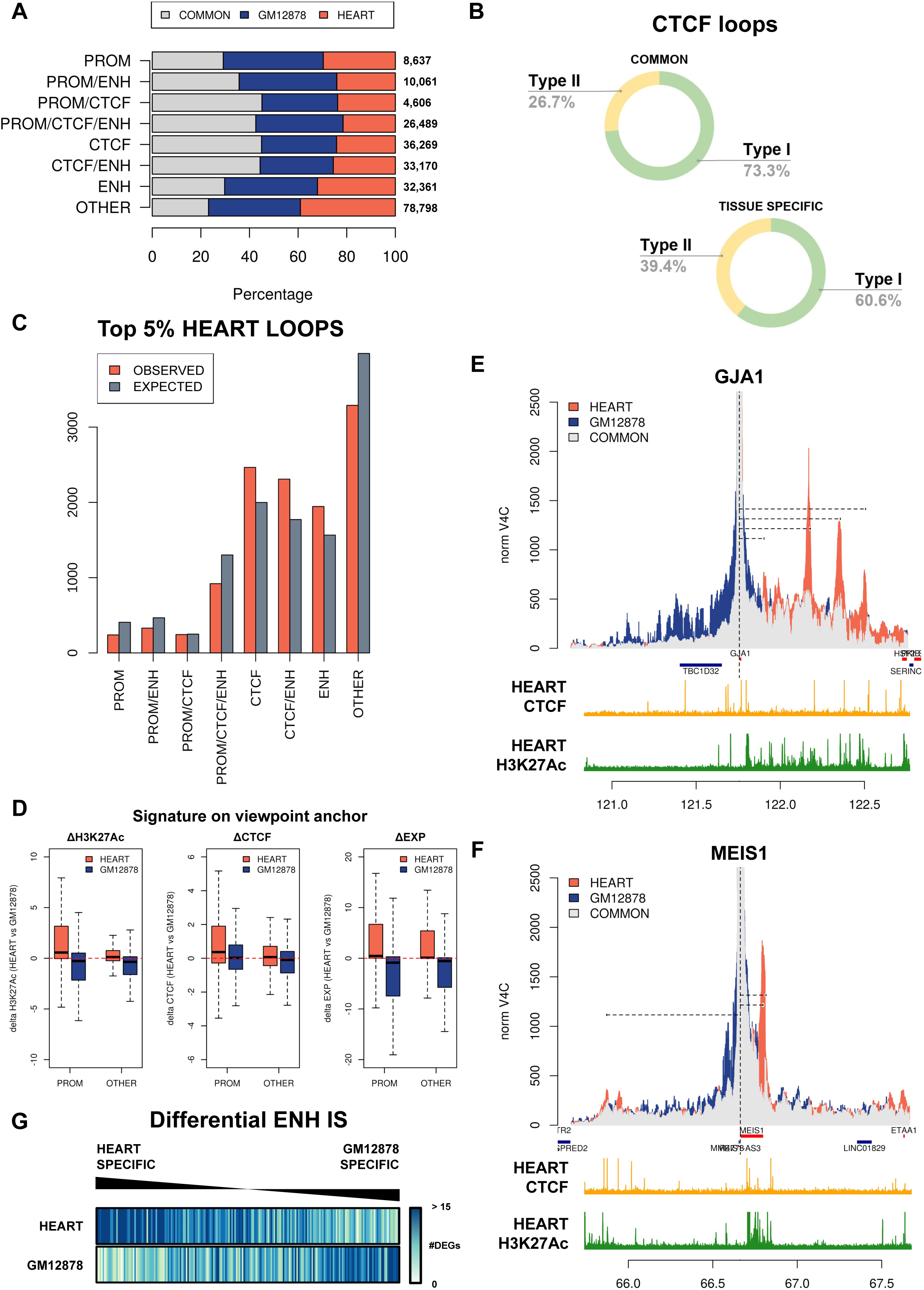
Tissue specific chromatin loops of the human heart. A) CTCF loops are most conserved (in relative and absolute terms) between heart and GM12878 cells. Plot shows per indicated category of peakHiC loops (as defined in heart) the proportion of shared and heart- and GM12878-specific chromatin loops. B) Class II CTCF loops are developmentally more dynamic than class I CTCF loops. Doughnut plots show that class II loops are relatively overrepresented among the tissue-specific loops (bottom) as compared to the shared loops (top). C) Each category of loops contributes proportionally to the top 5% most heart-specific loops (based on differential heart versus GM12878 interaction score). Plots shows per category the total number of observed versus expected loops among the top 5% most heart-specific loops. D) Tissue-specific loop formation coincides with tissue-specific accumulation of H3K27Ac (left) and CTCF (middle) at the anchors, and with increased expression of the nearest gene (right). Boxplots show the delta H3K27Ac ChIP-seq signal and delta CTCF ChIP-seq signal in heart over GM12878 at the loop anchors, and delta gene expression (RNA-seq RPKM) of the gene closest to the center of the anchor, for the top 5% heart tissue-specific loops (red) and top 5% GM12878 tissue specific loops (orange). E,F) Normalized V4C overlay contact profiles measured in heart (red) and GM12878 (yellow), of gene viewpoint GJA1 (E) and MEIS1 (F). In light grey are contacts shared between the tissues. Positive strand genes are indicated in red, negative strand genes in blue. Below each panel is shown the CTCF and H3K27Ac ChIP-seq results. G) Genome-wide quantitative correlation between tissue-specific differences in enhancer interaction scores and tissue-specific differences in transcriptional output. Genes are ranked to have those with the highest heart-specific enhancer interaction score on the left and those with the highest GM12878-specific enhancer interaction score on the right. Color code indicates the differential expression level of a gene (identified by using DESeq2), with blue being over-expressed and white being under-expressed in a given tissue (heart on top, GM12878 at bottom).

We then asked whether tissue-specific chromatin loop formation was associated with dynamic epigenetic alterations or transcriptional changes between the tissues. For this analysis, we quantitatively compared differential loop interaction scores to tissue-specific differences in their anchor-associated RNA-seq expression levels and CTCF- and H3K27Ac ChIP-seq signals. We focused on the 5% most tissue-specific chromatin loops and divided them into promoter-centered and non-promoter-centered loops (Figure 5D). The most heart-specific promoter loops showed heart-specific accumulation of H3K27Ac, both to the promoter anchor (Figure 5D) and to its interacting anchor (Figure S5D). We also found a clear heart-specific increase in expression of the genes engaged in these loops (Figure 5D). CTCF binding to these heart-specific loop anchors was similar between heart and GM12878 cells. Very similar results were seen for the category of non-promoter centered heart-specific chromatin loop: overall their anchors also showed higher levels of H3K27Ac, but not of CTCF, in heart than in GM12878 cells. Also, the expression of their closest gene was clearly higher in heart (Figure 5D). Corresponding epigenetic and transcriptional changes were observed for the 5% most GM12878-specific chromatin loops (Figure 5D). In a separate analysis, we further divided the 5% most heart- and GM12878-specific loops into the eight categories of loops described earlier, and analyzed their associated changes in H3K27Ac, CTCF and expression levels. As expected, results were very similar, but by separately considering CTCF-anchored loops it also became appreciable that tissue-specific accumulation of CTCF can accompany tissue-specific loop formation (Figure S5C).

We then looked further into the genes engaged in the 1% most prominent heart-specific loops. **Table 3** list these 200 genes, among which many well-established cardiac genes including the transcription factors *MEIS1*, *GATA4*, *GATA6*, and *TBX5*, as well as the genes encoding the connexin proteins *GJA1* and *GJA5.* Overlays of V4C contact profiles derived from heart and GM12878 Hi-C datasets highlight their strong heart-specific long-range chromatin interactions (Figures 5E **and** 5F). An intersection with pathways and traits defined by the Kyoto Encyclopedia of Genes and Genomes (KEGG) (Kanehisa et al. 2017) revealed an enrichment for cardiomyopathy traits, the PI3K-Akt, and Hippo signaling pathways among the genes involved in the top 1% most prominent heart-specific loops (Figure S5E). We then asked whether differential enhancer interaction scores were predictive of the differential expression levels of genes. Figure 5G ranks all genes according to their differential enhancer interaction score being highest either in heart (left) or in GM12878 (right) and shows their differences in expression levels between heart and GM12878 in color coding. The panels show that this ranking of genes based on their contact dynamics with enhancers clearly selects the genes that are most specifically expressed in GM12878 (top) and heart (bottom) (Figure 5G).

Collectively, the data shows that in particular tissue-specific enhancer activity and to a lesser extent tissue-specific CTCF binding events contribute to the strengthening of tissue-specific chromatin loops. Genes displaying major differences in enhancer contact frequencies between tissues typically also show corresponding changes in gene expression. The fact that the most dynamic non-promoter centered loops also often localize near tissue-specific genes suggests they may be relevant for their expression as well.

### Integration of cardiac specific chromatin loops with GWAS for cardiovascular traits and diseases

Most disease-associated loci identified in population based genetic studies or GWAS are located in non-coding regions of the genome and are presumed to modulate distal regulatory elements that loop to control the expression of target genes in disease-relevant tissue. Following this, we hypothesized that genetic variants associated with cardio-vascular traits should be enriched in the anchors of heart-specific chromatin loops and that the identification of these loops should help prioritizing the target genes. To investigate this, we first intersected the top 10% most specific chromatin loops of the GM12878 cell line and of primary cardiac tissue with genetic loci associated with human diseases and traits from the GWAS catalog. We found that there was a larger overlap of GWAS loci for cardiovascular (CVD, 14%) compared to non-CVD traits (8.6%) in cardiac tissue (Figure 6A). We then examined whether there was enrichment in the overlap between genetic loci associated with specific CVD or non-CVD diseases or traits and the top 10% strongest loops in GM12878 or cardiac tissue. In cardiac tissue, we observed strongest enrichment for loci involved in heart-related traits including PR interval (fold enrichment (FE_cardiac_) = 1.77, *p* = 4.3×10^-03^), atrial fibrillation (FE_cardiac_ = 1.51, *p* = 6.6×10^-03^) and the combined cardiovascular group (FE_cardiac_ = 1.33, p = 4.1×10^-05^), with nominal significance for PR and AF and significance after correcting for multiple testing for the combined cardiovascular group. For GM12878 loops, we observed Alzheimer’s (FE_GM12878_ = 2.51, *p* = 9.7×10^-04^), inflammatory bowel disease (FE_GM12878_ = 2.2, *p* = 7.4×10^-15^) and LDL (FE_GM12878_ = 2.04, *p* = 1.3×10^-03^) showing the strongest fold enrichment and significance after correcting for multiple testing (Figure 6B, Table S2). To further examine the relationship between the PR interval, atrial fibrillation, coronary artery disease, and the QT interval at the genome-wide level, we then plotted the summary level results for each disease or trait. In these modified Manhattan plots, we have illustrated the overlap of disease-associated loci with P-value < 5×10^-8^ and the top 10% strongest loops that are found in cardiac tissue or GM12878, showing that at many loci there is clearly more overlap with heart-specific loop anchors than with GM129878-specific loop anchors (Figure 6C, Figure S6).

**Figure 6.**
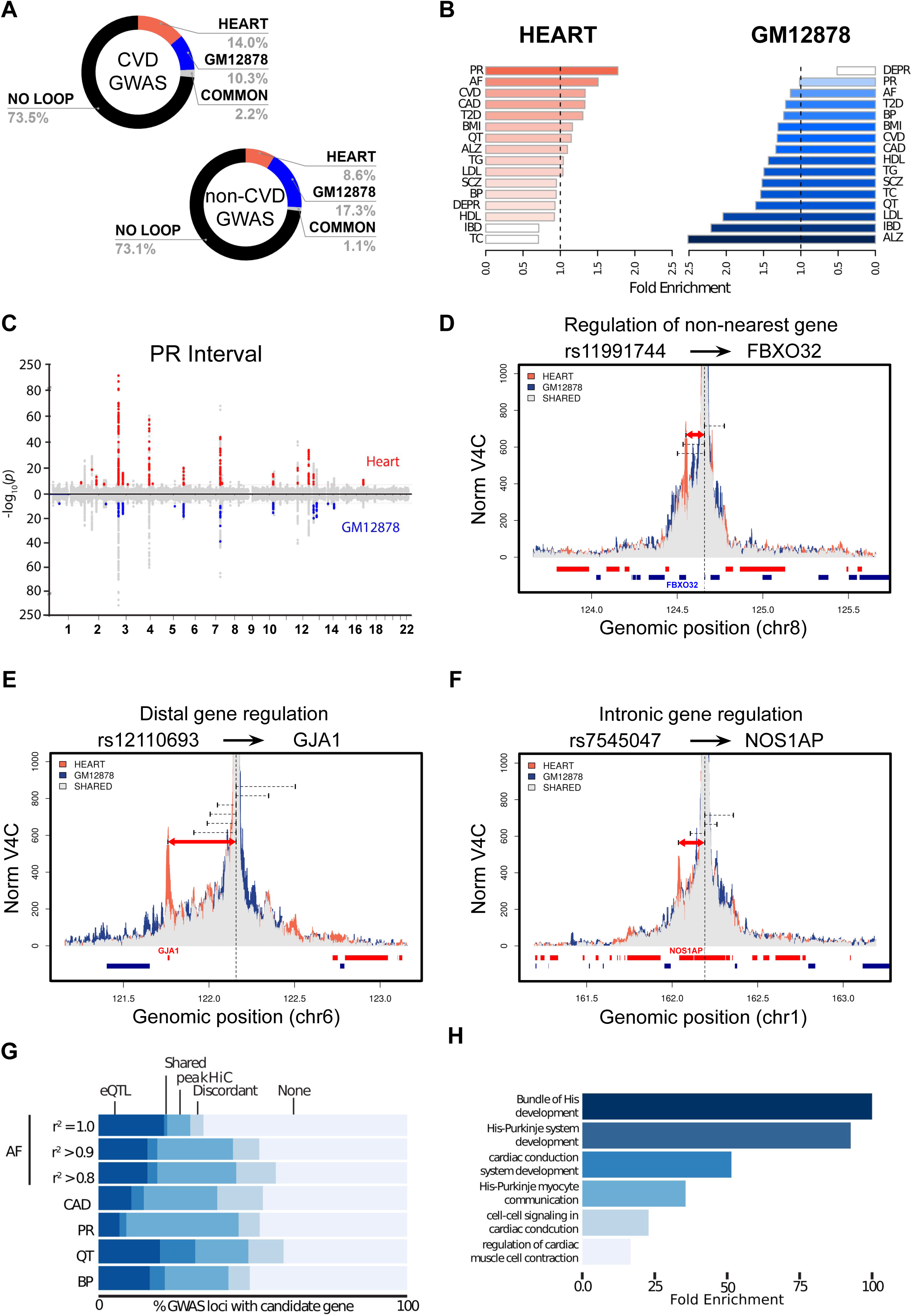
Relation between peakHiC data in human cardiac tissue and cardiovascular diseases. A) Doughnut plot detailing the proportion of sentinel variants at GWAS loci from the GWAS catalog with P-value < 5 x 10^-8^, intersecting with the top 10% strongest loops in GM12878 and/or heart. CVD: cardiovascular traits, non-CVD: non-cardiovascular traits (see **Table S3**). B) Results of enrichment analysis of top 10% strongest HiC loops in Heart and GM12878 for sentinel variants of various GWAS by GREGOR. AF: Atrial fibrillation, ALZ: Alzheimer’s, BMI: Body mass index, BP: Blood pressure, CAD: Coronary artery disease, CVD: combined cardio-vascular traits, defined in the methods, DEPR: Depression, IBD: Inflammatory bowel disease, LDL: LDL cholesterol, PR: PR interval, QT: QT interval, SCZ: Schizophrenia, T2D: Type 2 diabetes, TC: Total cholesterol, TG: Triglycerides. (*) indicates statistical significance. C) Manhattan plot for the PR interval from the electrocardiogram intersected with top 10% strongest peakHiC loop calls from heart (top) and GM12878 (lower, mirrored). Subthreshold (P-value > 5.0 x 10^-8^) SNPs were not labeled when overlapping with a loop. D) Virtual 4C plot demonstrating the looping of the heart rate-associated variant, rs1210693 with the promoter of GJA1. For all V4C plots in this figure, blue and red bars indicate genes on the + and – strands, respectively. Pink bars within the V4C histogram represent heart contact frequency, while orange bars represent the same in the GM12878 cell line. Gray colored bars indicate regions where the contact frequencies in both sample sources overlap. E) Virtual 4C plot demonstrating the looping of the QT interval-associated variant, rs7545047 with the promoter of GJA1. F) Virtual 4C plot demonstrating the looping of the PR interval-associated variant, rs11991744 with the promoter of FBXO32. G) Bar graph detailing the percentage of loci for which a candidate gene was observed using eQTL analyses, peakHiC, or both. R2 refers to the SNPs considered around the top variant for the various AF loci. For CAD, PR, QT and BP an R2 of 0.8 was utilized. BP: blood pressure. H) Bar graph detailing the results of gene ontology analysis using Panther. Results were generated from genes identified as looping to GWAS loci in heart tissue for the following electrophysiological diseases/traits: atrial fibrillation, PR interval, and QT interval.

We then examined the local V4C profiles at a series of GWAS loci, to identify a number of interesting categories of cardiac specific loops linked to disease loci (exemplar plots in Figure 6D-6F **and** Figure S7). One category is displayed in Figure 6D in which an intragenic disease SNP loops to a distant gene. rs11991744 is a sub-threshold locus for the PR interval, is intronic to *KLHL38*, yet loops to *FBXO32,* an E3 ubiquitin ligase implicated in muscle atrophy. Another example of an intragenic variant that based on Hi-C may well be involved in the regulation of another, more distal, gene is rs17608766, which is associated with blood pressure (P = 2 x 10^-46^) (Kichaev et al., 2019) and QRS interval duration (P = 1 x 10^-44^) (Prins et al., 2018a) and is located intronic to GOSR2, yet loops to *WNT9B* (Figure S7A).

A second category highlights risk variants located in large gene deserts that form strong heart-specific loops over large distances to a specific flanking gene. This is illustrated by the *GJA1* locus. The SNP rs12110693 has been associated with heart rate (P = 1.6 x 10^-9^)(Cho et al., 2009), resides in a 1Mb region devoid of any genes, and loops in a cardiac specific manner to the *GJA1* gene encoding the gap junction protein connexin 43 (Figure 6E). Similar examples were also observed with variants associated with sudden cardiac death (Aouizerat et al., 2011) (rs4621553, P = 4.12 x 10^-8^) and the ion channel *KCNN2* as well as the PR interval (Verweij et al., 2014) (rs10850409, P = 7.62 x 10^-13^) and the cardiac specific transcription factor *TBX3* (Figure S7B **and** S7C).

A more common scenario is an intragenic disease associated SNP looping to the promoter of the same gene. This pattern is illustrated by rs7545047 located in the genetic locus most significantly associated with the QT interval (Arking et al., 2014). This variant is intronic to *NOS1AP* and loops to the proximal promoter of the *NOS1AP* gene (Figure 6F). Similar results were observed at the rs652889 locus for the QT interval (P = 8 x 10^-7^) (Marroni et al., 2009), which is located inside the large *PTPRG* gene and loops over more than 200kb to its promoter, and the PR interval associated region (rs267567, P = 4.14 x 10^-11^) (Butler et al., 2012) which locates inside the *ITGA9* gene and loops to its promoter (Figure S7D **and** 7E).

With these patterns in mind, we then sought to quantify the additive benefit of using Hi-C in combination with peakHiC to facilitate the identification of candidate genes at GWAS loci. We began by using a large GWAS of AF as a reference dataset and determined how many AF genetic loci looped to the TSS of a candidate gene in the region. By considering variants in strong LD with the lead SNP (defined as a region bounded by an R^2^ ≥ 0.8), we were able to identify candidate genes for many AF associated loci, an approach that was then extended to coronary disease and blood pressure as well as electrocardiographic traits. Traditionally, expression quantitative trait loci mapping (eQTL) is used to link a disease associated variant to transcription of candidate genes at the locus (Figure 6F **and Table S3**). We found that using peakHiC identifies candidate genes for many additional loci over eQTL mapping alone (Figure 6G). Using candidate genes identified for cardiac conduction diseases and traits (atrial fibrillation, PR interval, and QT interval), we then performed biological pathway enrichment analysis using Panther (Figure 6H **and** Table S4) where we identified significant enrichment for a range of cardiac conduction related biological processes.

## Discussion

### Hi-C to link disease variants to genes

More than 90% of disease-associated genetic variation uncovered by GWAS lies in non-coding DNA. Such variants do not change the functionality of a gene product but instead are thought to affect transcript levels through distant gene regulation. In support of this concept, many of these genomic sites display transcription regulatory potential in disease-relevant cell-types, as judged from their epigenetic signatures (1000 Genomes Project Consortium et al., 2010; Chen et al., 2016; Maurano et al., 2012). Gene regulation over distance requires chromatin looping to bring distal regulatory DNA elements in close proximity to target genes. In order to nominate target genes of non-coding risk variants it is therefore instrumental to uncover their long-range chromatin interactions. Since regulatory DNA elements act and can form chromatin loops in a cell type-specific manner, it is best to study contacts involving disease loci in the disease-relevant cell type.

Hi-C is the most unbiased and complete genome-wide strategy to investigate spatial chromatin conformations and is therefore in principle best suited to identify physical contacts between and among genes, regulatory DNA elements and genetic risk variants. It is not trivial though to identify chromatin loops in a sensitive and meaningful manner from Hi-C datasets, even with very high sequencing depth. For that reason, researchers currently use alternative, more targeted chromosome conformation capture strategies such as Hi-ChIP (Mumbach et al., 2017) and promoter-capture Hi-C (PCHi-C) (Javierre et al., 2016; Martin et al., 2015) to topologically connect GWAS variants to genes. Hi-ChIP analyzes proximity ligation junctions after chromatin pulldown with an antibody against a selected chromatin factor or modification, to target loop analysis exclusively to genomic sites carrying this factor(Mumbach et al., 2017). Hi-ChIP results are therefore influenced by differences in protein occupancy and thus pulldown efficiencies between genomic sites and the method cannot uncover chromatin loops between anchors not carrying the selected factor or modification. PCHi-C is insensitive to protein occupancy issues but ignores non-promoter anchored chromatin loops and fails to provide contact profiles of promoter-interacting regions (PIRs), for reciprocal loop confirmation. PCHi-C has been used to prioritize candidate target genes of GWAS hits associated with autoimmune diseases and blood cell traits (Burren et al., 2017; Cornish et al., 2019; Javierre et al., 2016; Martin et al., 2015), colorectal cancer (Jäger et al., 2015; Orlando et al., 2018), breast cancer (Baxter et al., 2018) and bone mineral density (Chesi et al., 2019). Two recent studies applied PCHi-C to link genes to GWAS hits associated with cardiovascular diseases (Choy et al., 2018b; Montefiori et al., 2018). In both studies, human in vitro induced pluripotent stem cells-derived cardiomyocytes (CM) were used instead of primary human donor heart tissue. CM contact libraries were sequenced at an estimated depth of >1.5B read pairs. Using CHiCAGO, a dedicated algorithm to extract loops from capture Hi-C datasets (Cairns et al., 2016), one study found 18,159 promoters involved in an average of 35 (maximum = 433) interactions (Choy et al., 2018a), while the other study identified 14,954 promoters engaged in an average of 27 (maximum = 757) interactions (Montefiori et al., 2018). For comparison, peakHiC identified 26,327 promoters being engaged in an average of 2.7 interactions (maximum = 25), plus ∼23,000 non-looped promoters in heart.

Every peak caller relies on thresholding, which in the case of Hi-C analyses will always be arbitrary since the method collects contact pairs from populations of cells that individually carry differently structured genomes (Finn et al., 2019; Krijger and De Laat, 2013). For every Hi-C identified contact pair one could argue it must be present in at least the tiniest fraction of cells. However, we recommend that our peakHiC criteria, being that loops must be (1) reproducibly found across Hi-C replicates, (2) confirmed individually by reciprocal peak calling, (3) confirmed collectively by APA, (4) visually appreciable from V4C contact maps and optionally (5) correlating with known looping associated features such as transcriptional output and epigenetic signatures, are essential for meaningful extraction of recurrently formed chromatin loops from Hi-C and Hi-C-related datasets. Frequent spatial proximity between distal genomic sites does not automatically imply a functional interaction: genomic contact data can therefore only help to prioritize candidate target genes of risk variants and regulatory DNA sequences. Since peakHiC more selectively filters recurrent chromatin interactions than PCHi-C and additionally employs Hi-C datasets that interrogate not just promoter but all DNA contacts in an unbiased and genome-wide manner, we propose that using Hi-C in combination with peakHiC is the preferred method to interpret GWAS hits in their 3D genomic context and nominate the molecular mechanisms they may influence to contribute to disease.

### Hi-C-derived V4C contact profiles to link individual genes to risk variants

Regardless of peak calling results, we emphasize the importance of visual inspection of overlay V4C contact profiles from different tissues, to best appreciate and prioritize the candidate relevant long-range contacts of an individual gene, regulatory sequence or risk variant. Extracting V4C plots from Hi-C data is not novel (Sexton et al., 2012; Zhang et al., 2012) and several online tools for the analysis and visualization of Hi-C data offer the possibility to extract and look at V4C contact profiles (Harmston et al., 2015; Robinson et al., 2018; Wang et al., 2018). Our V4C profiles differ in that we compile the contacts of a few dozen neighboring DpnII restriction fragments centered on a sequence of interest (a gene promoter, risk variant, CTCF site, regulatory DNA element) and then use a sliding running mean window to plot their contacts. This results in highly reproducible contact profiles: overlay V4C plots from the heart left atrium and ventricle are nearly always indistinguishable and even GM12878 and heart V4C plots most often look similar overall, with tissue-specific long-range chromatin contacts only to be found at particular sites of interest. This overall reproducibility and our demonstration that tissue-specific chromatin contacts generally relate to tissue-specific transcription patterns and protein binding events, highlights the importance to search for obvious cell-type specific long-range contacts, which is best done by inspection of overlays of V4C contact profiles from different cell types. Binning contacts comes with a loss in resolution, but at 10kb resolution it is often possible to nominate the gene promoter, CTCF site or regulatory DNA element responsible for loop formation.

### PeakHiC uncovers large-scale contact networks with regulatory potential

With peakHiC we uncovered several interesting chromosome topology features. Overall, our data demonstrate that a much larger percentage of CTCF sites than anticipated are involved in chromatin looping. These may have been previously missed possibly because they occur in smaller percentages of cells, hence giving weaker signals in population-based Hi-C contact matrices. We identify a class of CTCF loops (intradomain class II loops) that generally do not adhere to the rule that productive loop formation requires interactions between convergently oriented CTCF bound sites (de Wit et al. 2015; Sanborn et al. 2015). We demonstrate by APA the validity of these loops. Intra-domain class II loops are relatively small and developmentally dynamic and often share an anchor with the larger domain (class I) loops that predominantly form between convergent CTCF binding sites. We and others recently showed that multiple distal CTCF sites can spatially aggregate (Allahyar et al., 2018; Bintu et al., 2018), which we proposed can be the consequence of collisions (‘traffic jams’) between chromatin extruding cohesin complexes. We speculate that class II loops between non-convergent CTCF sites may also be formed as a consequence of loop collision events. We further observe that looping density, not only between genes and regulatory sequences but also between CTCF binding sites, is much higher in the A than B compartment. This supports the idea that active loop formation may be required to maintain an open chromatin configuration (Schwarzer et al., 2017). We find extensive contact networks among multiple gene promoters (as seen before (Dao et al., 2017; Li et al., 2012b)), between promoters and shared enhancers (also frequently observed in PCHI-C experiments, see e.g. (Choy et al., 2018a)), and between protein-coding gene promoters and lncRNA promoters (Engreitz et al., 2016). Interestingly, we also frequently observe long-range contacts between genes and sites with a repressive chromatin signature. In all cases interactions correlate with the expected corresponding transcriptional output. An important advantage of peakHiC is its capacity to assign interaction scores to all long-range chromatin contacts. This now enables ranking contacts based on their loop strength, to prioritize for each category the loops that are most interesting for future systematic functional validation studies. Using a distance-normalized interaction score we were able to show that loop strength correlates with CTCF, cohesin and H3K27Ac association strength, as measured by ChIP-seq. This quantitative relationship not only supports a causal relationship between these factors and loop formation but also underscores that both Hi-C and ChIP measurements (semi-)quantitatively reflect cell population frequencies of (DNA and protein) contact events.

### Connecting CVD risk variants to genes in the heart 3D genome

We initiated these studies to topologically connect cardiovascular disease associated variants in non-coding DNA to possible target genes. The rate of disease-associated loci discovery has far outpaced the ability of the field to identify the gene targets and mechanisms underlying the loci (reviewed in Gallagher and Chen-Plotkin, 2018). The most simplistic method, merely assigning the most proximal gene as the target, has a sound basis when considering the results of mammalian enhancer trap studies (Ruf et al., 2011), but is rife with the potential for false positives in the absence of additional information. Differential expression analyses in the heart through expression quantitative trait loci (eQTL) measurements have improved our target assignment (GTEx Consortium et al., 2013; Roselli et al., 2018), but are limited by sample numbers, the ephemeral nature of RNA species, and modest effect sizes for even the starkest examples. In short, the effect on gene expression, at least in bulk tissue analyses, is often too subtle to be reliably detected for all loci. Clearly, additive approaches are necessary to improve gene target prediction form GWAS loci; one of which is presented herein.

By applying peakHiC to primary human heart Hi-C data we were able to link 151 CVD GWAS loci (R^2^ >0.9) to distal genes. In aggregate, this is a large increase in the number of loci for which we have a plausible gene target in the relevant tissue type when compared to eQTL studies alone. The results for peakHiC-called genes were particularly encouraging in the combined analysis of cardiac electrophysiology traits where, in addition to known genes implicated in cardiac electrophysiology such as *GJA5* (Christophersen et al., 2013; Gollob et al., 2006) and *TBX5* (Ma et al., 2016; Nadadur et al., 2016), a global examination of biological processes associated with the genes identified yielded GO terms centered on the signal conduction and contraction of the myocardium.

The results of eQTL analyses and peakHiC are rarely overlapping, a result likely due to the limitations of each approach. The former relies upon strong effect sizes, where proximal regulation is enriched. However, due to the resolution limits of peakHiC, and all 3C methods, these contacts are difficult to successfully detect due to the decay curves generated from the ligation of nearby regions in 2-dimensional space. However, eQTL methods rarely detect distal gene regulation, a class of events particular suited to chromatin capture methods. Instead of proposing the superiority of one approach, we suggest that the methodologies are excellent complements to each other, aiding in gene identification for loci where the other method may be weak. Importantly, neither eQTL nor peakHiC gene target prioritization is sufficient to conclusively implicate a gene in disease or trait. Follow up studies on gene function using relevant biological models, whether cell or model organism based, should be employed to convincingly establish a role for this regulatory event in the given trait.

## Acknowledgments

This work was supported by a Fondation Leducq (14CVD01) Transatlantic Network grant to Drs. de Laat and Ellinor, and by grants from the National Institutes of Health to Dr. Tucker (1K01HL140187), Dr. Margulies (R01HL105993) and to Dr. Ellinor (1RO1HL092577, R01HL128914, K24HL105780). This work was also supported by a grant from the American Heart Association to Dr. Ellinor (18SFRN34110082) and an AHA SFRN postdoctoral fellowship to Dr. Hall. This research was also supported by the Carol and Roch Hillenbrand and George L. Nardi, MD, funds at Massachusetts General Hospital. This work was also supported by funding from the Oncode Institute to Dr. de Laat.

## Author Contributions

Conceptualization: VB, GG, WdL

Methodology: VB, GG, WdL

Software: VB, GG

Formal Analysis: VB, GG, CR

Investigation: CREH, VB, GG, CR, NT

Resources: MCH, JFM, KBM

Data Curation: VB, GG

Writing - Original Draft: VB, GG, WdL, NT, PTE

Writing - Review & Editing: VB, GG, WdL, NT, CR, AH, PTE, KBM

Visualization, VB, GG, NT, CR

Supervision: WdL, PTE

Project Administration: WdL, PTE

Funding Acquisition: WdL, PTE

## STAR Methods

### Contact for Reagent and Resource Sharing

Further information and requests for resources and reagents should be directed to and will be fulfilled by the Lead Contacts, Patrick T. Ellinor or Wouter de Laat.

### Experimental Model and Subject Details

*Human Subjects:* Adult human heart tissue samples were provided by The Myocardial Applied Genomics Network (MAGNet; www.med.upenn.edu/magnet) at the University of Pennsylvania. Heart were obtained from brain-dead organ donors free of any structural heart disease after next of kin provided written informed consent for research use of donated tissue. Tissue use for research was approved by the relevant institutional review board at the Gift-Of-Life Donor Program, Philadelphia, PA.

### Method Details

#### Hi-C data

Paired-end Hi-C sequencing reads were aligned independently with bwa using its BWA-MEM algorithm with default parameters to the hg19 reference genome. Reads with both mates mapped were further processed and filtered using the HiCUP (Wingett et al. 2015) pipeline to filter out experimental artifacts (including non-digested and religated fragments) and PCR duplicates. The resulting valid Hi-C contact pairs from left atrium (LA), left ventricle (LV) and GM12878 cell line were associated respectively to DpnII, DpnII and MboI restriction fragments, and used for downstream processing and analysis.

#### PeakHiC: definition of virtual 4C viewpoints and contact profiles

In order to transform Hi-C data into viewpoint centered Virtual 4C (V4C) profiles, we first associated each 1bp candidate viewpoint to its residing MboI/DpnII fragment. Given the sparsity of Hi-C data, we need to pool captures across a number of restriction fragments neighbouring the intended viewpoint fragment in order to increase the amount of captures available for analysis. We choose to pool 31 fragments for each V4C viewpoint (see below and **Table 2** for justification). We then count the number of Hi-C contact pairs between the pool of 31 viewpoint fragments and all MboI/DpnII restriction fragments within a 2Mb interval (1Mb upstream and 1Mb downstream, including intra-viewpoint contact pairs). The total number of Hi-C contacts, for each viewpoint, was normalized by scaling with a constant to 1 million Hi-C contact pairs per analyzed V4C profile (2 Mb) to aid visual comparison between profiles. Finally, V4C contact profiles were computed by calculating a running mean of normalized fragment coverage across 31 consecutive restriction fragments.

#### PeakHiC: peak calling – first round

For the lymphoblastoid cell line (GM12878), we used 9 different published MboI digested in-situ Hi-C libraries as input to peakHiC (Rao et al., 2014). For human heart Hi-C from left atrium (LA) and left ventricle (LV), we used the data from 4 independent Hi-C templates from different donors described in this manuscript. We adapted peakC (Geeven et al., 2018) to identify peaks in Hi-C data using the replicate V4C profiles in these 3 different Hi-C datasets. peakC is a robust non-parametric method developed initially for the identification of contact peaks in 4C-Seq and Capture-C data. Contact peaks are genomic regions that in replicate experiments show a statistically significant increase in contact frequency over the expected background. peakC fits a background model independently for every replicate V4C profile using monotonic (isotonic) regression and uses rank-product based statistics to identify fragments that score significantly above background in replicate experiments. It requires a significance threshold parameter alphaFDR which aims to control the expected proportion of false positives at the desired level and an additional threshold qWr (ratio with respect to background) parameter for further effect size filtering. We ran the algorithm with parameters set to alphaFDR=0.05, qWr=1. We called peaks from 493,937 ‘CTCF viewpoints’ and 49,052 ‘TSS’ viewpoints described in the main text. Loops that thus met peakC significance thresholds were further filtered to be at least 30Kb apart.

#### PeakHiC: peak calling – reciprocality condition

Loop anchors from peaks identified by peakHiC from viewpoints in the first round are kept only if they are found reciprocally, i.e. any loop anchor from a V4C viewpoint x must contain (another) V4C viewpoint which makes a loop to a 10Kb loop anchor region that contains x. To determine this, all loop anchors identified in the first round of loop calling are first re-sized to 10Kb regions centered at the maxV4Cscore (Figure 2H) position. In case a resized loop anchor did not contain any V4C viewpoints from the first round, we could not immediately verify the reciprocality condition. Instead we took all these ‘orphan’ loop anchors as V4C viewpoints in a second round of peak calling and kept only those loops for the re-sized 10Kb loop anchor regions centered at the maxV4Cscore position contained the original V4C viewpoint (from the first round).

#### PeakHiC: loop interaction score calculation

For robust coverage-rank based scoring of Hi-C loop interactions across different V4C viewpoints and different Hi-C maps (from different cell types / conditions) we did the following. PeakHiC loop anchor summits (maxV4CScorePos, Figure 2H) are defined as the genomic position (bp) of the MboI / DpnII fragment where the maximum peakC V4C score is attained. For each peakHiC peak, we resize both the viewpoint region and its contacting anchor to bins of 10kb. For the viewpoint region, we center the 10 kb bin at the site of interest (TSS, CTCF binding site, etc), for the contacting anchor we center the 10Kb bin at the maxV4CScorePos. For each Hi-C library (replicate / condition) we counted the number of valid Hi-C contacts between these pairs of 10Kb regions (one pair of regions for each peakHiC loop). We next summed the Hi-C contacts across different replicates to get the Hi-C loop interaction score (IS) for each loop in GM12878, LA, LV and HEART (Hi-C libraries from LA and LV combined). We then rank each peakHiC loop within each Hi-C map based on its IS and compute its coverage rank quantile (a number between 0 and 1) relative to all other loops (covQ). Loops in the top and bottom 2.5% based on coverage (covQ) were filtered out.

#### Total enhancer interaction score calculation

Per gene, a total enhancer interaction score (totENH-IS) is computed. We selected all the loops having genes as viewpoints and contacting loop anchors overlapping with enhancers as identified with MACS on ChIPseq data from H3K27Ac (paragraph “ChIP-seq data analysis”). Per gene we finally summed up the interactions scores (IS) from each loop on H3K27Ac peaks.

#### Loop distance normalization

To intersect Hi-C and ChIP-seq data and analyze whether there is a relationship between loop interaction score (influenced by loop distance) and protein occupancy at the looping anchors (not influenced by loop distance) it is necessary to first normalize the loop interaction score for distance. To normalize for the distance between both anchors of a peakHiC loop on Hi-C loop coverage, we binned loops based on this distance in 150 bins, such that each bin contains an equal number of loops. Next, we determined the average loop interaction score (covQ values) of all loops in a bin and subtracted this bin specific average. The resulting covQ.norm value is used for distance normalized interaction score ranking of peakHiC loops. We used this interaction score measure in all instances where Hi-C loop strength (distance dependent) was compared to ChIP-seq protein occupancy levels (not distance dependent). (Dixon et al., 2012)

#### Creating a non-redundant set of loops

We collected all the peakHiC reciprocal loops from HEART (LA+LV) and GM12878 for a total collection of 722,816 loops, by using ∼540,000 TSS and CTCF viewpoints. For each loop we have two coordinates referring to the viewpoint position (VP) and the genomic position of the summit of the interaction peak (maximum V4C score). We denoted with X and Y the most up- and downstream positions of each loop. Redundancy is removed in the following way:

– Genome is partitioned in 10Kb sized bins and a unique identifier is assigned to each bin;
– For each loop we assign two unique identifiers, as those from the residing bin for the anchor X and Y. Any loop with the same pair of unique identifiers has both the anchors in the same pair of 10Kb genomic bins;
– If multiple loops occur in the same pair of genomic bins, we keep only the loop with highest read coverage in its 10Kb anchors.

#### PeakHiC: V4C viewpoint size determination

Selection of an appropriate V4C viewpoint size for peakHiC is a tradeoff between resolution and coverage/reproducibility. In order to evaluate the minimum number of fragments to pool and be used for the (meta) viewpoint region, we tested 5 different ‘fragment pool’ sizes of 11, 21, 31, 41 and 51 MboI/DpnII fragments centered on the fragment of interest. For these candidate viewpoint sizes we generated V4C profiles for a collection of 4756 gene promoters viewpoints located on chr1, thus generating 4756 profiles per candidate viewpoint size. For each V4C profile and for all different viewpoint sizes we then computed a series of metrics, reported in **Table 2**, in the format mean(sd):

– the average viewpoint region size in bp (VPsize);
– the 10th and 90th percentile of the viewpoint region size;
– the average number of Hi-C contact pairs, per replica as well as for the total collection of replicates, in a 0.2Mb and 2Mb region surrounding the viewpoint excluding those mapping on the viewpoint region itself;
– the average coverage, namely the percentage of fragments covered by at least one Hi-C contact pair, per replica as well as for the total collection of replicates, in a 0.2Mb and 2Mb region surrounding the viewpoint.

Based on our previously defined criteria to judge whether a 4C experiment is of sufficient quality for meaningful interpretation (Van De Werken et al., 2012), we required to have a minimum local coverage of 40% (0.2 Mb around the viewpoint). Following that requirement, we binned 31 fragments per viewpoint, giving, per replicate Hi-C experiment, an average of 545 contacts and 46% coverage in the 0.2 Mb interval (total across all replicates: 4357 contacts and 85% coverage) (**Table 2**).

#### A and B compartments identification

A and B compartments in the heart Hi-C datasets (LA and LV) have been identified by using the *Eigenvector* function on the Hi-C datasets from juicer tools (Durant et al. 2016); the sign of the eigenvector indicates the compartment. For the GM12878 Hi-C data, we used the A and B compartment data as previously published (Rao et al. 2014).

#### Insulation score and directionality index calculation

We implemented in R the insulation score calculation as defined by Olivares-Chauvet et al. 2016. Scores were calculated in regions +/- 250Kb at 2Kb resolution. In heatmaps, visualized scores were multiplied by −1 and mean zero normalized such that high positive scores reflect strong contact insulation. For the directionality index (DI) calculation we used the definition by (Dixon et al., 2012). DI scores were also calculated at 2Kb resolution. Heatmaps display z-score normalized values of the DI.

#### Classification of CTCF loops

For classification of CTCF loops into type I and type II we first quantified the total bias in directionality index (for each region centered around a looped CTCF site) by taking the difference of the directionality index averaged over the 50 (2Kb spaced) bins up- and downstream of the CTCF site. We then computed a running mean statistic to calculate enrichment in directionality bias for the CTCF sites ordered by CTCF ChIP-Seq enrichment (Figure 2D, Figure 4B). This statistic averages the directionality bias at each site over the 1000 CTCF sites nearest in the ranking and we determined the threshold at the first CTCF site for which this running average went below 0. All CTCF sites above this threshold are classified as type I, with strong directionality bias and high insulation average, and other CTCF sites are classified as type II.

#### ChIP-seq data analysis

ChIP-Seq NGS paired end reads were aligned to the hg19 human genome with Bowtie2 (REF) using default settings, and duplicated reads were removed. Peaks were called using the MACS software (v2.1.0) (Zhang et al., 2008) with default parameters. Peak enrichment was determined for each peak identified by MACS2 using R with Bioconductor (Gentleman et al., 2004) and compEpiTools (Kishore et al., 2015) packages. In short, the enrichment is defined as log2(PeakSig—inputSig), where PeakSig is the reads count on the enriched region in the ChIP sample, inputSig is the reads count on the same region in the corresponding input sample, both normalized by millions of mapped reads.

#### RNA-seq data analysis

RNA-seq NGS paired end reads were aligned to the hg19 human genome using the TopHat aligner (version 2.1.0)(doi: 10.1186/gb-2013-14-4-r36) with default parameters and duplicated reads were removed. Reads count associated to each gene based on the UCSC-derived hg19 GTF gene annotation were generated using the featureCounts function using R with Bioconductor and Rsubread package (doi: 10.1093/nar/gkz114).

Gene expression was defined by determining reads per kilobase per million of mapped reads (RPKM) and by defining the total library size as the number of reads mapping to exons only. Differentially expressed genes were identified using the Bioconductor package DESeq2 (Love et al., 2014) and by imposing q-value lower than 0.01 and absolute log2 Fold Change > 0. For each gene we selected the most active TSS by selecting the one with the highest amount of RNA-seq mapped reads in the region +/- 1Kb around each TSS.

#### KEGG pathway analysis

KEGG pathway enrichment analysis has been computed on the genes found in the top 1% HEART specific loop anchors, after sorting them by differential normalized interaction score HEART vs GM12878. A total of 443 genes have been provided to ConsensusPathDB (Kamburov et al., 2013) with default parameters and plotting has been performed by selecting the top 5 ranked KEGG pathways.

### Functional and GWAS annotation

#### HiC loops for intersection

We ordered the HiC loops (N=230,391) by diff.CovQ.norm and selected the top 10% loops (N=23,039) with the strongest diff.CovQ.norm value for Heart (left ventricle and left atrium) and GM12878 respectively. We included the viewpoint and anchor regions for each HiC loop for the intersections.

#### Overlap of CVD versus non-CVD GWAS loci with HiC loops

We identified sentinel variants for CVD loci from the GWAS catalog (Buniello et al., 2019) as well as additional sentinel variants for AF (Christophersen et al., 2017a) and P-wave indices (Christophersen et al., 2017b). Included traits for the CVD group were: Brugada syndrome, P terminal force, P wave duration, PR interval, PR segment, QRS complex, QRS duration, QT interval, TPE interval measurement, atrial fibrillation, cardiac hypertrophy, cardiovascular disease - type II diabetes mellitus, cardiovascular measurement, cardiovascular measurement - left ventricular hypertrophy, congenital heart disease, congenital heart malformation, congenital left-sided heart lesions, dilated cardiomyopathy, heart failure, heart function measurement, heart rate and sudden cardiac arrest. For the non CVD group we kept the remaining traits in the GWAS catalog excluding the previously listed CVD traits as well as: cardiovascular disease, systolic blood pressure, diastolic blood pressure, blood pressure, coronary artery calcification, abdominal aortic artery calcification, hypertension, myocardial infarction, coronary heart disease, atherosclerosis, resting heart rate, RR interval, coronary heart disease - myocardial infarction, coronary artery bypass - atrial fibrillation, coronary artery bypass - vein graft stenosis, cardiac arrhythmia, coronary artery bypass - myocardial infarction, heart failure - mortality, response to statin - coronary heart disease, stroke - coronary heart disease, large artery stroke - coronary heart disease, Conotruncal heart malformations - parental genotype effect measurement, Conotruncal heart malformations, large artery stroke - coronary heart disease, large artery stroke, carotid artery intima media thickness, coronary artery bypass - Acute kidney injury - serum creatinine measurement and coronary artery bypass - myocardial infarction. For each group we selected sentinel variants with P-value < 5×10-8 keeping the variant with the lowest P-value within a 1mb window for each trait. Across traits we identified independent sentinel variants, keeping variants with the lowest P-value and removing all correlated variants with r2 > 0.8 based on European ancestry LD from 1000 Genomes phase3 v5 (1000 Genomes Project Consortium et al., 2015). We intersected the sentinel variants from the CVD and non-CVD group with the top 10% strongest HiC loops in heart tissue and in the GM12878 cell line.

#### Enrichment

We calculated the enrichment using the tool GREGOR v1.4.0 (Schmidt et al., 2015) for reported sentinel variants for the complex diseases: atrial fibrillation (AF) (Roselli et al., 2018), alzheimer’s (ALZ) (Lambert et al., 2013), coronary artery disease (CAD) (van der Harst and Verweij, 2018), depression (DEPR) (Wray et al., 2018), inflammatory bowel disease (IBD) (de Lange et al., 2017), schizophrenia (SCZ) (Schizophrenia Working Group of the Psychiatric Genomics Consortium, 2014) and type 2 diabetes (T2D) (Scott et al., 2017), as well as the traits: body mass index (BMI) (Locke et al., 2015; Speliotes et al., 2010) blood pressure (BP) (Warren et al., 2017), PR-interval (PR) (van Setten et al., 2018), QT-interval (QT) (Arking et al., 2014), high density cholesterol (HDL) (Willer et al., 2013), low density cholesterol (LDL) (Willer et al., 2013), triglycerides (TG) (Willer et al., 2013) and total cholesterol (TC) (Willer et al., 2013). We also created the group combined cardiovascular (CVD) including sentinel variants from the traits and diseases: AF, CAD, BP, PR, QT, heart failure (Aragam et al., 2018), heart rate (den Hoed et al., 2013) and QRS-duration (Prins et al., 2018b), echocardiographic traits of diastolic function, LV structure and systolic function (Wild et al., 2017). We used the following settings to calculate proxies: r2 threshold of 0.8, a window size of 500kb and European ancestry. The size of the neighborhood in GREGOR was defined as 1,000. The enrichment was calculated for the top 10% strongest HiC loops in heart tissue and in the GM12878 cell line. Bonferroni corrected significance threshold was established as P-value < 1.56×10-3.

#### Manhattan plots

We generated mirrored Manhattan plots of the summary results of the traits AF, PR and QT. Highlighted are genome-wide significant variants (P-value < 5×10-8) that intersect with the top 10% strongest HiC loops in heart tissue (red, highlighted in the upper half) and GM12878 (orange, highlighted in the lower half). We used the (R Core Team, 2015) package qqman (Turner, 2018) for plotting.

#### GWAS-to-gene identification

For all sentinel variants from the traits AF, PR, QT and CAD we identified the proxies with r2 > 0.8, based on linkage disequilibrium (LD) from a European reference set from the Broad AF study (Roselli et al., 2018) calculated in PLINK v1.90 (Chang et al., 2015). We identified the nearest gene to the sentinel variant using the RefSeq gene reference. We annotated the sentinel variants and proxies with genes identified via expression quantitative trait loci (eQTL) from human LA samples from the Myocardial Applied Genomics Network repository (Chang et al., 2015), as well as eQTLs from the Genotype-Tissue Expression (GTEx) project (GTEx Consortium, 2015) in the cardiac tissues right atrial (RA) and left ventricular (LV). Additionally, we annotated genes where the anchor region of any called peak-HiC loops in LA and LV overlapped with the gene’s TSS.

#### Gene set enrichment

To display the enrichment of heart loops in biologically relevant pathways that are specific to CVD, we used the Gene Ontologies (Ashburner et al., 2000; The Gene Ontology Consortium, 2017) and performed gene set enrichment on loops which intersected with a SNP or proxy (r2 > 0.8) for the following traits: AF, BP, CAD, PR, QT. We used PantherDB (Thomas et al., 2006) to determine enrichment for the GO biological process. For the first 10 terms, we plotted the number of genes present, and the raw P-value, as no terms were significant due to the small number of genes per trait plotted. For combining the PR and QT gene lists, a short list of pathways were significantly enriched by FDR correction, and for that figure the FDR corrected and adjusted P-values are displayed.

#### Data and Software Availability

##### Data Access

The Hi-C data from the human heart and GM12878 cell line have been linked to genetic variation for a range of cardiovascular diseases and traits using the peakHiC algorithm. This data will be made publicly available upon publication of the manuscript and can be found at the Broad Cardiovascular Disease Knowledge Portal (www.broadcvdi.org). The human sequence data generated in this study are available at dbGaP (https://www.ncbi.nlm.nih.gov/gap/), accession number phs001539.v1.p1.

##### Software

The algorithm from peakHiC is written in R (https://www.R-project.org/) and is available on GitHub at the following link: https://github.com/vbianchi/peakHiC.

## Tables

**Table 1:** Left atrium (LA) and left ventricle (LV) Hi-C mapping report.

**Table 2:** Statistical analysis and report of different Virtual 4C viewpoint bin sizes.

**Table 3:** Top 200 genes associated to top 1% most dynamic heart loops.

## Supplemental Information

**Figure S1.**
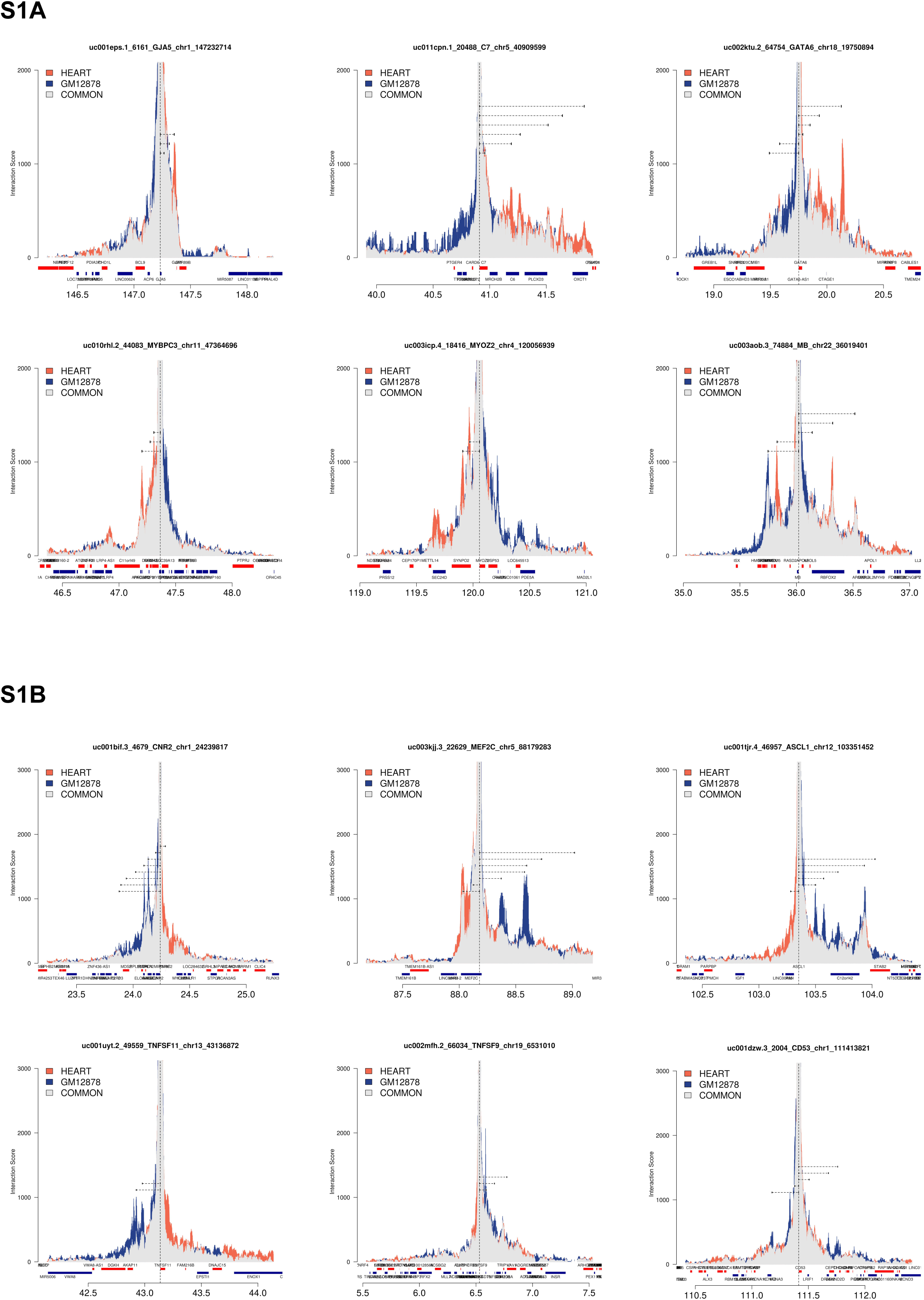
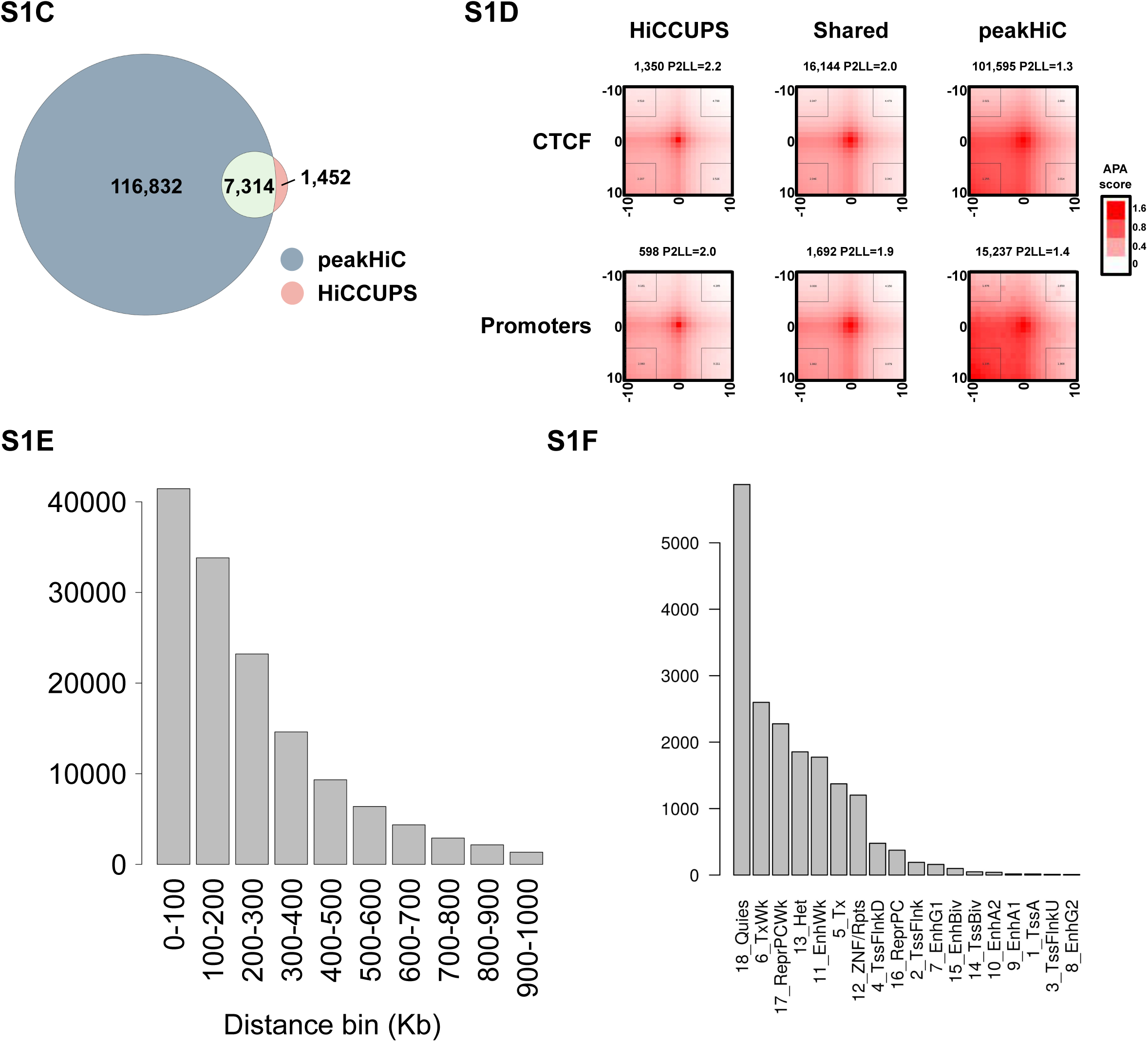
**A)** Normalized V4C overlay contact profiles showing the local (-/+ 1 Mb) contacts of GJA5, C7, GATA6, MYBPC3, MYOZ2 and MB gene promoters. Contact frequencies found both in HEART and GM12878 are in grey, contact frequencies unique to HEART or GM12878 are in red and blue, respectively. The viewpoint is marked by vertical black dashed lines, vertical dashed lines show peakHiC-called loops. Below each V4C plot: positive strand genes in red, negative strand genes in blue. **B)** Normalized V4C overlay contact profiles showing the local (-/+ 1 Mb) contacts of CNR2, MEF2C, ASCL1, TNFSF11, TNFSF9 and CD53 gene promoters. Contact frequencies found both in HEART and GM12878 are in grey, contact frequencies unique to HEART or GM12878 are in red and blue, respectively. The viewpoint is marked by vertical black dashed lines, vertical dashed lines show peakHiC-called loops. Below each V4C plot: positive strand genes in red, negative strand genes in blue. **C)** Venn diagram showing the amount of chromatin loops identified in GM12878 cell line by peakHiC (blue), HiCCUPS (pink) and from both methods (green). **D)** Heatmap plots that show the result of aggregate peak analysis (APA) which measures normalized contact enrichment of Hi-C contacts between pairs of regions. The six panels show contact enrichment for CTCF and promoter centered loops identified in GM12878 cell line by HiCCUPS only, peakHiC only and both methods (Shared). **E)** Barplot showing the distribution of genomic loops identified with peakHiC at different distance bins from the viewpoint, ranging from 0 to 1000Kb with increment of 100Kb. **F)** 18-state ChromHMM categories representation from GM12878, at the loop anchors of loops whose viewpoint anchor do not overlap any CTCF peak, H3K27Ac peak and gene promoters (OTHER category in Figure 1E) descendant sorted by frequency.

**Figure S2.**
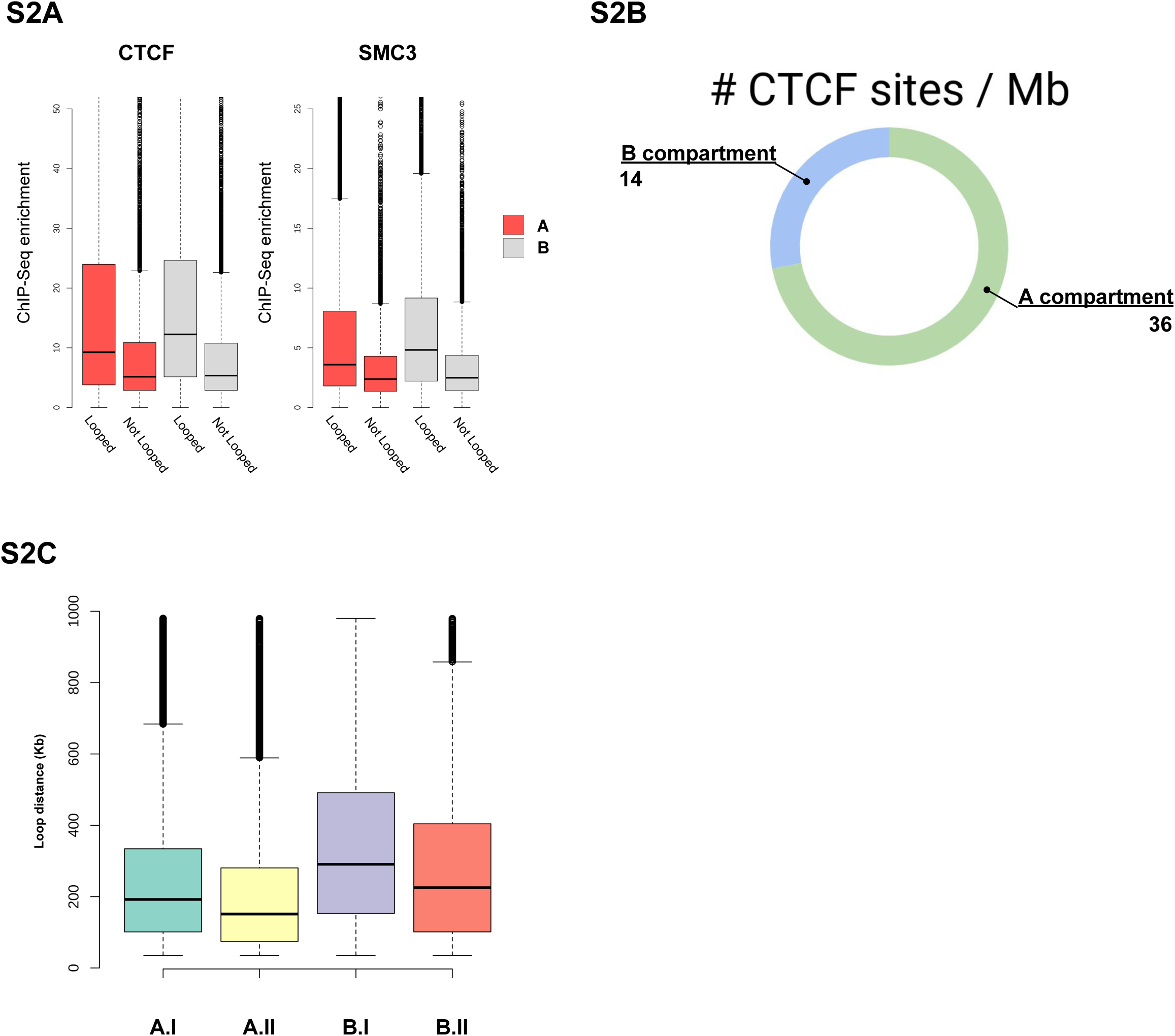
**A)** Boxplots that show the distributions of CTCF and SMC3 ChIP-Seq enrichment at CTCF sites that are / are not looping, split according to their location in the A or B compartment in GM12878 cells. **B)** Doughnut plot showing the density of CTCF bound sites in A and B compartment in GM12878 cell line. **C)** Boxplots that show the distributions of loop distances (between VP and loop anchor) for the sets of loops defined in Figure panel 2C.

**Figure S3.**
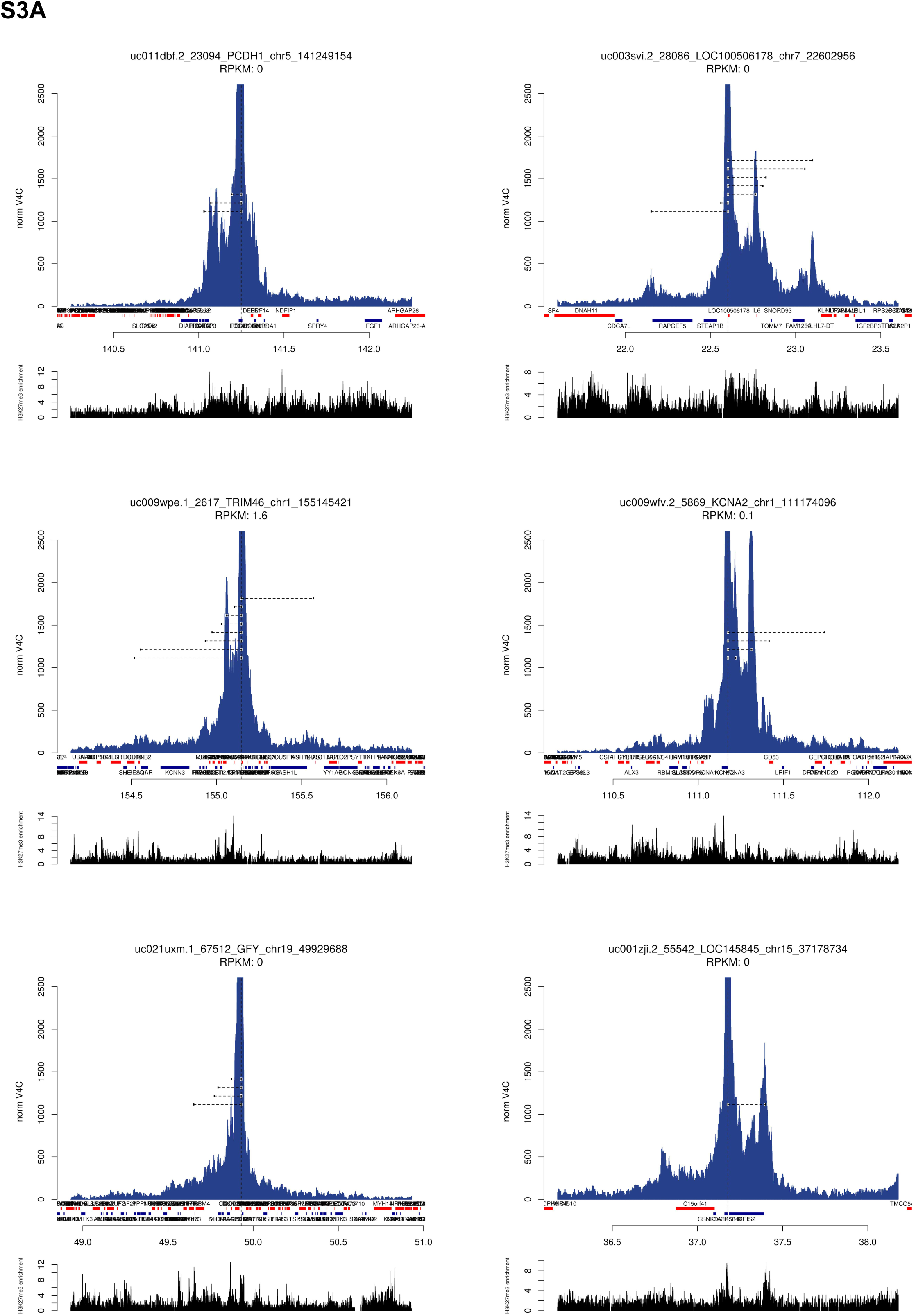
**A)** V4C contact profiles in GM12878 for gene promoters PCDH1, LOC100506178, TRIM46, KCNA2, TMIE and LOC145845. The viewpoints are marked by vertical black dashed lines, positive strand genes in red, negative strand genes in blue. peakHiC loops identified on top of repressor regions, identified by H3K27me3 ChIP-seq mark in GM12878, are marked with black boxes (10Kb in size) connected by an horizontal dash line.

**Figure S4.**
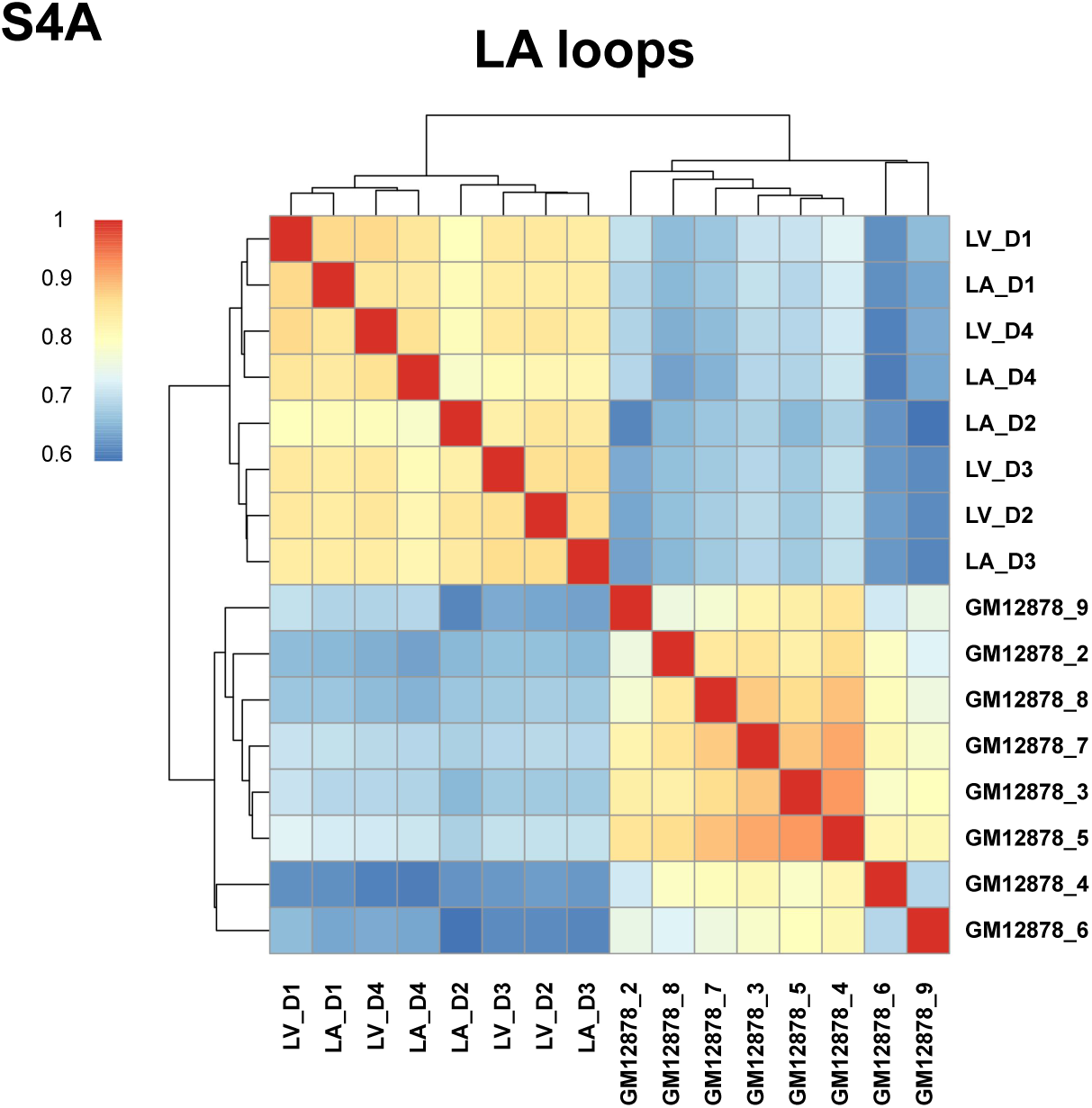
**A)** Heatmap plots that show the correlation in Hi-C contact counts between the 10Kb anchors of peakHi-C loops in LA samples. Spearman’s rank correlation was computed for all independent Hi-C templates in LV (4 reps), LA (4 reps) and GM12878 (8 reps).

**Figure S5.**
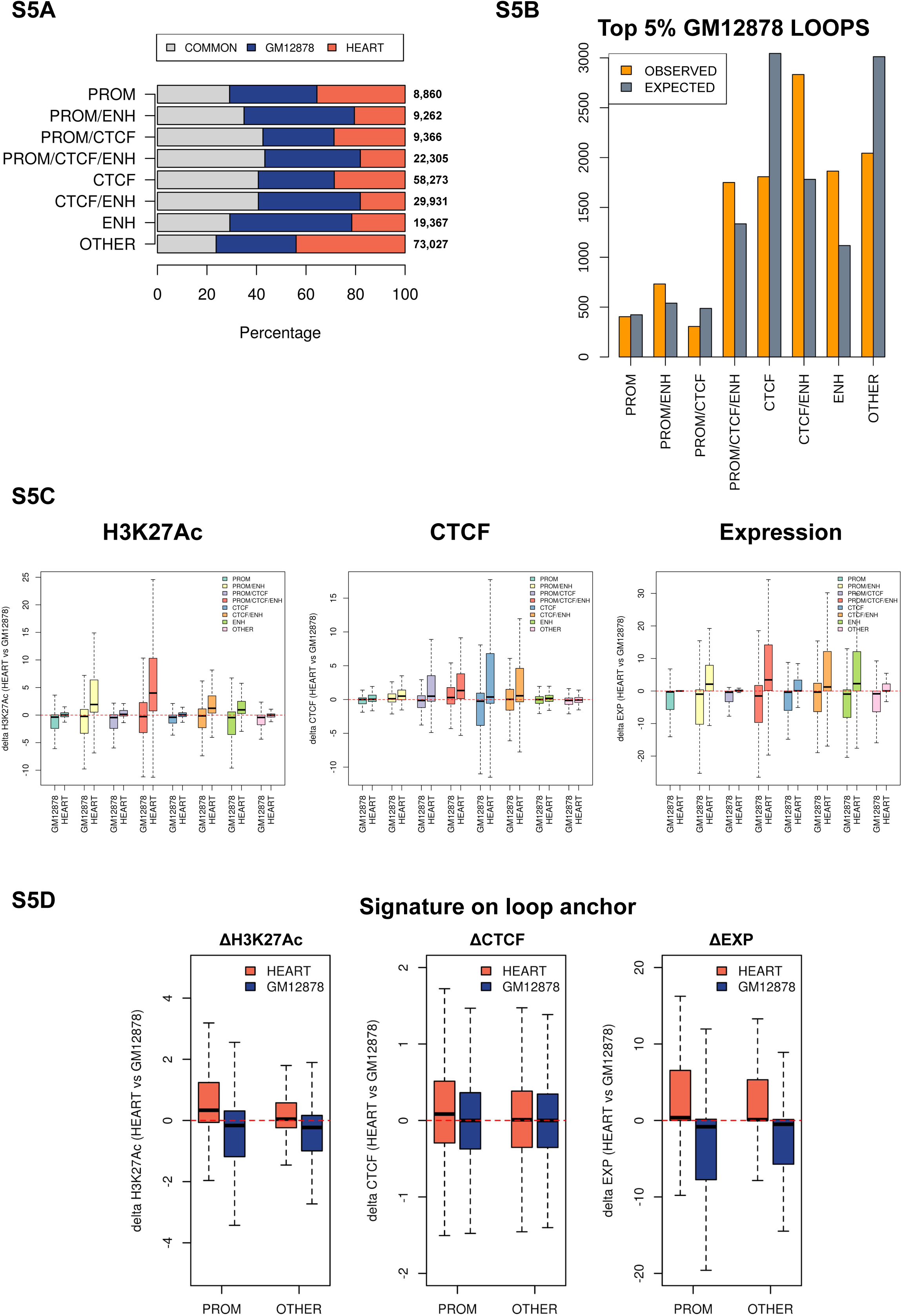
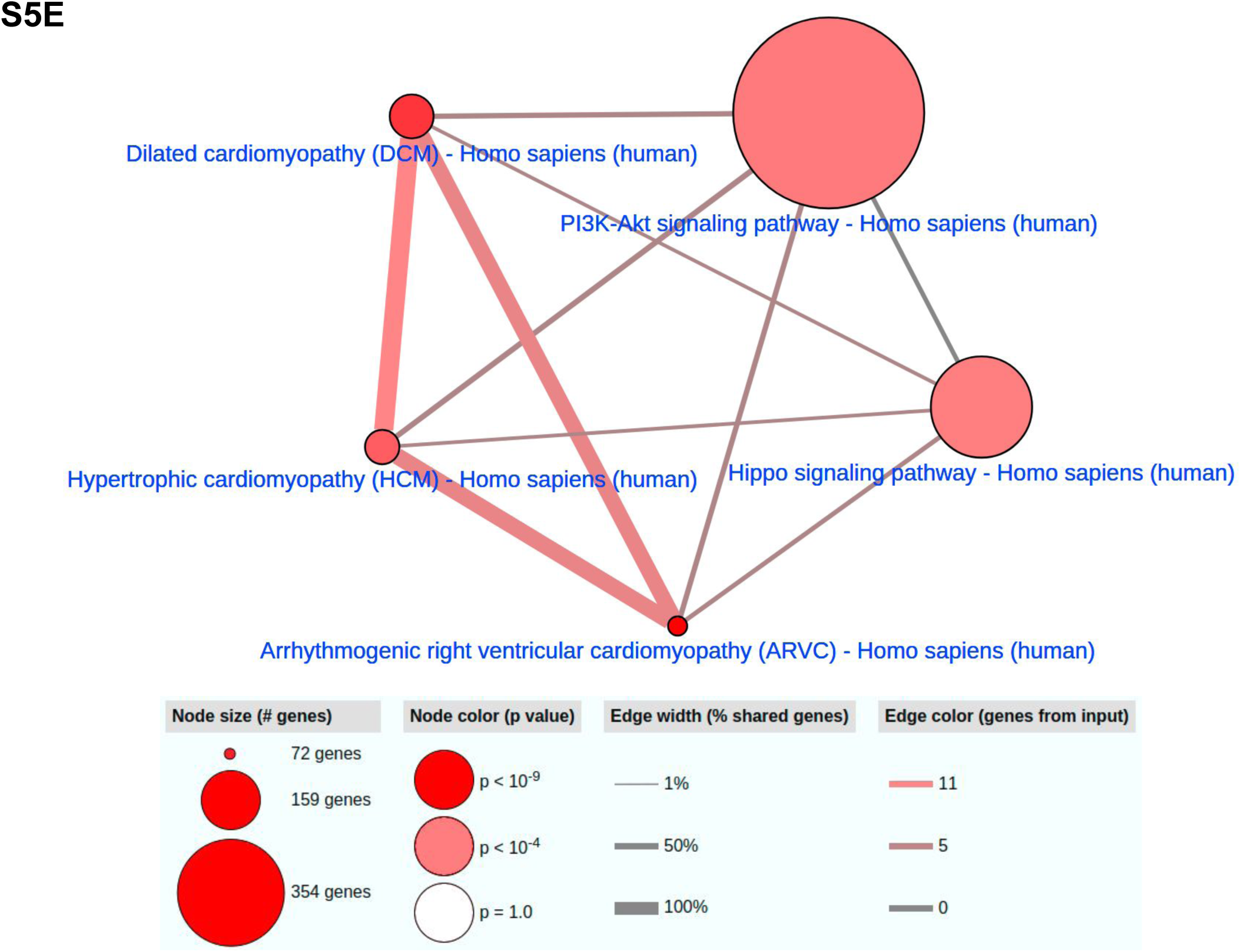
**A)** Stacked barplot showing the amount of peakHiC loops in common and tissue specific in HEART vs GM12878 cell line. Category have been created by having the 10Kb viewpoint anchor overlapping GM12878 CTCF ChiPseq peaks (CTCF), GM12878 H3K27Ac peaks (ENH), gene promoters (PROM), any possible configuration of these three marks (CTCF/ENH, PROM/CTCF/ENH, PROM/ENH, PROM/CTCF), or none of the above (OTHER). **B)** Barplot showing the distribution per category identified in Figure S5A of the top 5% loops GM12878 specific vs expected. Loops have been selected after sorting them by differential normalized interaction score GM12878 vs HEART (LA+LV) cell line. **C)** Series of boxplots showing the delta H3K27Ac (ChIP-seq signal), delta CTCF (ChIP-seq signal) and delta gene expression (RNA-seq RPKM of the closest gene to the center of the anchor) HEART vs GM12878 at the anchors of top 5% HEART tissue specific loops (red) and GM12878 tissue specific loops (blue). **D)** Series of boxplots showing the delta H3K27Ac (ChIP-seq signal), delta CTCF (ChIP-seq signal) and delta gene expression (RNA-seq RPKM of the closest gene to the center of the anchor) HEART vs GM12878 at the loop anchors of top 5% HEART tissue specific loops and GM12878 tissue specific loops, categorized by having the 10Kb viewpoint anchor overlapping GM12878 CTCF ChiPseq peaks (CTCF), GM12878 H3K27Ac peaks (ENH), gene promoters (PROM), any possible configuration of these three marks (CTCF/ENH, PROM/CTCF/ENH, PROM/ENH, PROM/CTCF), or none of the above (OTHER). **E)** KEGG pathway enrichment analysis computed on the genes found in the top 1% HEART specific loop anchors. These loops have been identified after sorting them by differential normalized interaction score HEART vs GM12878.

**Figure S6.**
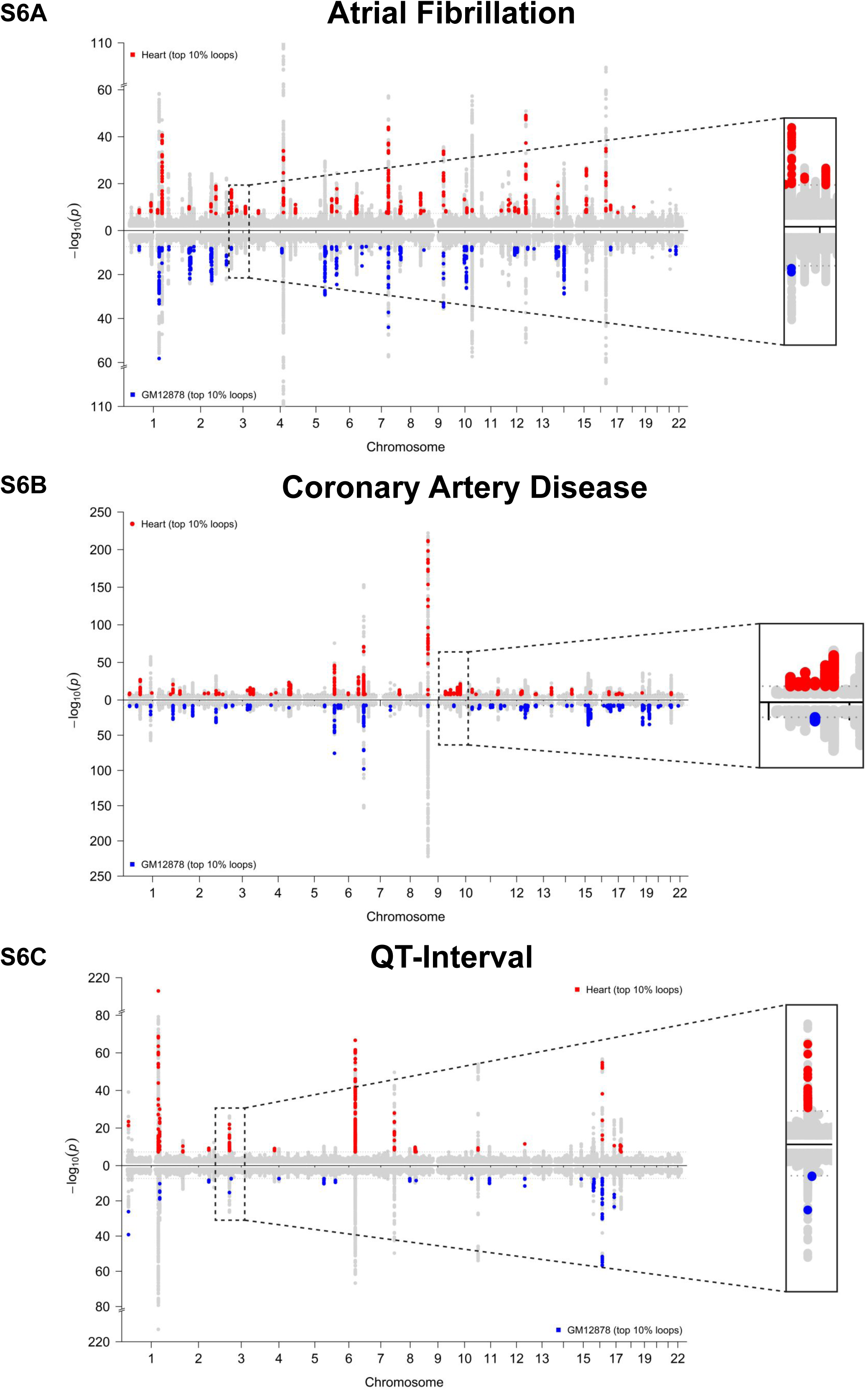
Related to Figure 6. Manhattan plots detailing overlap between TSS-centered loops and GWAS SNPs. Manhattan plots for A) atrial fibrillation, B) coronary artery disease, and C) the QT interval of the electrocardiogram intersected with the top 10% strongest peakHiC loops from heart (red) or GM12878 (blue). Subthreshold (P-value > 5.0 x 10-8) SNPs were not labeled when overlapping with a loop. Insets for all three plots detail regions where top variants at GWAS loci were only found to intersect with heart chromatin loops.

**Figure S7.**
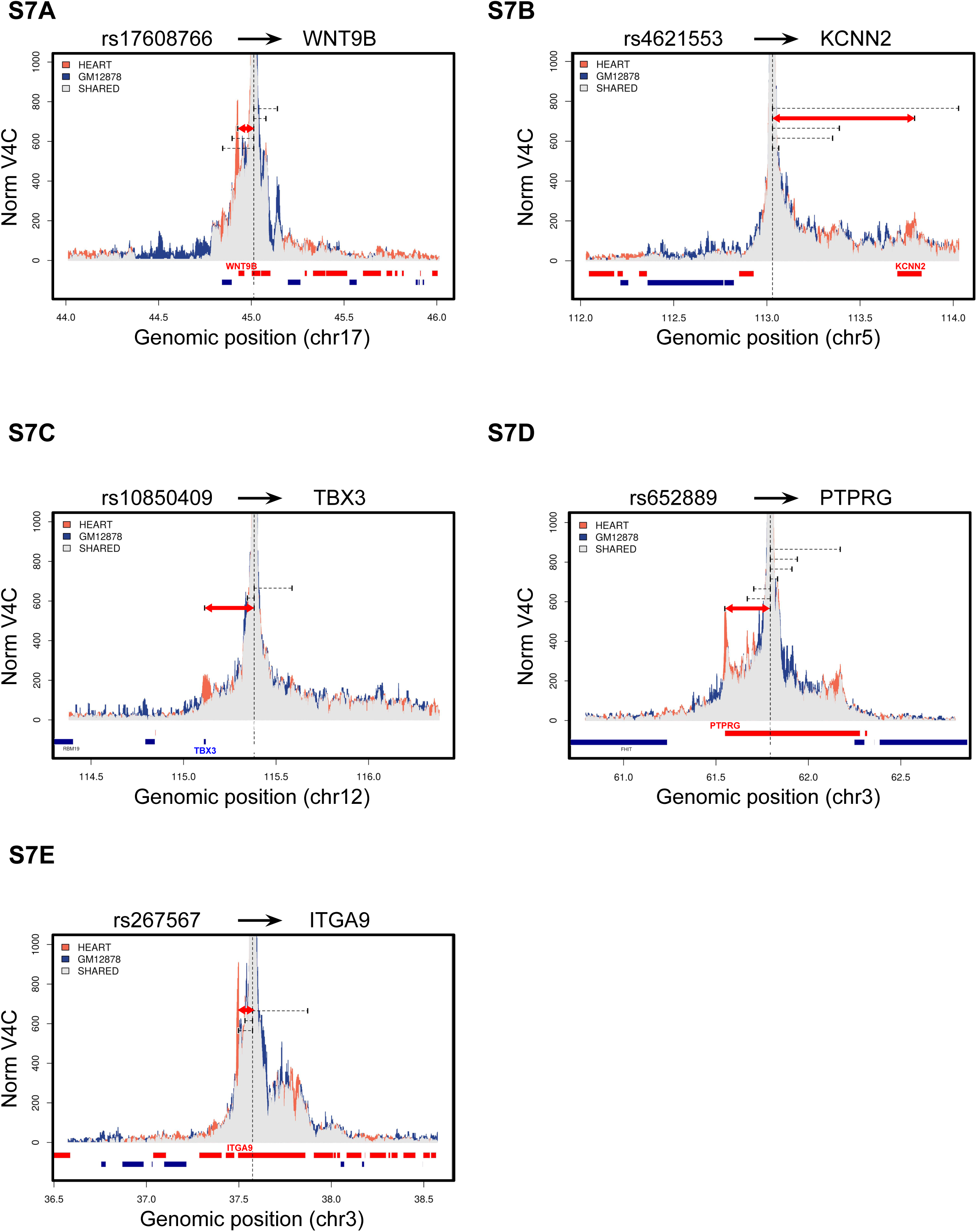
Related to Figure 6C-F. Exemplar virtual 4C profiles at GWAS loci.

**Table S1.** Baseline characteristics of human heart donors

| Sex | Age (years) | Race | Height (cm) | Weight (kg) | Heart mass (g) | HMI (g/m<sup>2</sup>) | LVEF (%) | Creatinine (mg/dl) |
| --- | --- | --- | --- | --- | --- | --- | --- | --- |
| Female | 24 | White | 160 | 75 | 262 | 143.50 | 61 | 0.6 |
| Female | 55 | White | 168 | 57 | 343 | 210.31 | 70 | 0.5 |
| Male | 53 | Black | 193 | 100 | 443 | 191.33 | N/A | 0.8 |
| Male | 43 | White | 173 | 102 | 432 | 195.12 | 60 | 1 |

HMI: Heart mass index, LVEDD: left ventricular end diastolic dimension, LVESD: left ventricular end systolic dimension, PW Thick: Posterior wall thickness, LVEF: Left ventricular ejection fraction.

**Table S2.**
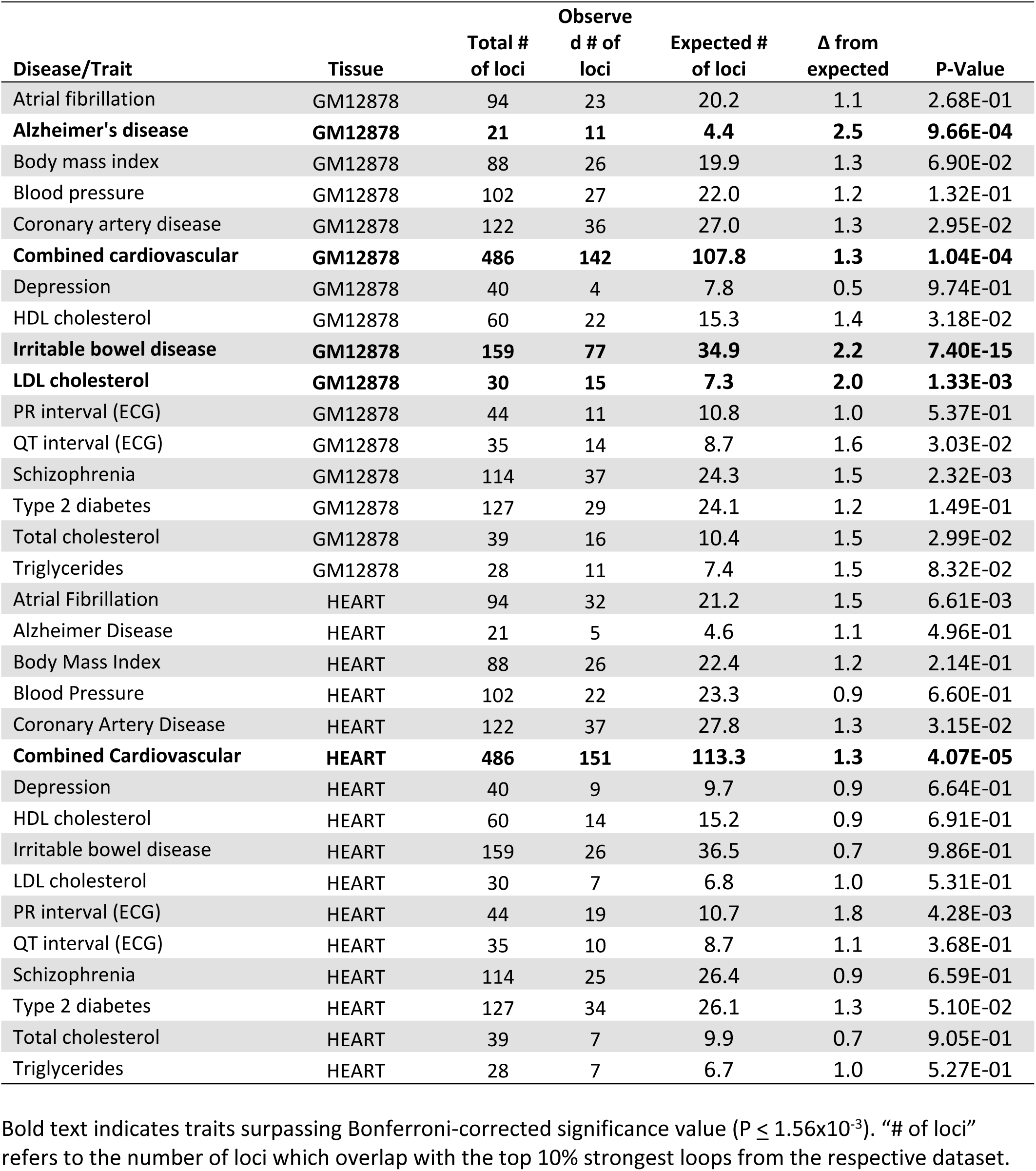
Results of global enrichment analysis using GREGOR

**Table S3.**Candidate genes identified for cardiovascular GWAS loci.

Please see the attached .csv table

**Table S4.** GO biological pathways enrichment analysis for electrophysiological GWAS loci by Panther.

| **GO biological process complete** | **Genes in**<br>**GO term** | **HiC genes**<br>**observed** | **Expected** | **Fold**<br>**enrichment** | **raw P-**<br>**value** | **FDR adjusted**<br>**P-value** |
| --- | --- | --- | --- | --- | --- | --- |
| bundle of His development (GO:0003166) | 4 | 3 | 0.03 | > 100 | 8.96E-06 | 1.42E-02 |
| His-Purkinje system development (GO:0003164) | 5 | 3 | 0.03 | 92.63 | 1.43E-05 | 1.50E-02 |
| cardiac conduction system development (GO:0003161) | 12 | 4 | 0.08 | 51.46 | 2.81E-06 | 6.36E-03 |
| bundle of His cell to Purkinje myocyte communication (GO:0086069) | 13 | 3 | 0.08 | 35.63 | 1.37E-04 | 4.43E-02 |
| cell-cell signaling involved in cardiac conduction (GO:0086019) | 27 | 4 | 0.17 | 22.87 | 4.51E-05 | 2.04E-02 |
| regulation of cardiac muscle cell contraction (GO:0086004) | 37 | 4 | 0.24 | 16.69 | 1.38E-04 | 4.37E-02 |
| ventricular cardiac muscle tissue development (GO:0003229) | 54 | 5 | 0.35 | 14.29 | 3.88E-05 | 2.28E-02 |
| regulation of striated muscle contraction (GO:0006942) | 93 | 8 | 0.6 | 13.28 | 2.92E-07 | 4.62E-03 |
| regulation of cardiac muscle contraction (GO:0055117) | 77 | 6 | 0.5 | 12.03 | 1.60E-05 | 1.58E-02 |
| ventricular septum development (GO:0003281) | 69 | 5 | 0.45 | 11.19 | 1.16E-04 | 4.36E-02 |
| positive regulation of muscle organ development (GO:0048636) | 70 | 5 | 0.45 | 11.03 | 1.23E-04 | 4.54E-02 |
| positive regulation of striated muscle tissue development (GO:0045844) | 70 | 5 | 0.45 | 11.03 | 1.23E-04 | 4.43E-02 |
| positive regulation of muscle tissue development (GO:1901863) | 71 | 5 | 0.46 | 10.87 | 1.31E-04 | 4.42E-02 |
| heart contraction (GO:0060047) | 87 | 6 | 0.56 | 10.65 | 3.06E-05 | 2.31E-02 |
| striated muscle contraction (GO:0006941) | 108 | 7 | 0.7 | 10.01 | 9.63E-06 | 1.27E-02 |
| cardiac ventricle development (GO:0003231) | 124 | 8 | 0.8 | 9.96 | 2.25E-06 | 8.91E-03 |
| action potential (GO:0001508) | 95 | 6 | 0.62 | 9.75 | 4.88E-05 | 2.15E-02 |
| heart process (GO:0003015) | 99 | 6 | 0.64 | 9.36 | 6.07E-05 | 2.53E-02 |
| regulation of muscle contraction (GO:0006937) | 162 | 9 | 1.05 | 8.58 | 1.67E-06 | 8.82E-03 |
| regulation of striated muscle tissue development (GO:0016202) | 134 | 7 | 0.87 | 8.06 | 3.61E-05 | 2.29E-02 |
| regulation of muscle tissue development (GO:1901861) | 137 | 7 | 0.89 | 7.89 | 4.14E-05 | 2.04E-02 |
| regulation of muscle organ development (GO:0048634) | 138 | 7 | 0.89 | 7.83 | 4.32E-05 | 2.07E-02 |
| cardiac muscle tissue development (GO:0048738) | 161 | 8 | 1.04 | 7.67 | 1.40E-05 | 1.58E-02 |
| cardiac chamber development (GO:0003205) | 165 | 8 | 1.07 | 7.49 | 1.66E-05 | 1.54E-02 |
| muscle cell development (GO:0055001) | 147 | 7 | 0.95 | 7.35 | 6.34E-05 | 2.57E-02 |
| regulation of muscle system process (GO:0090257) | 234 | 10 | 1.52 | 6.6 | 4.17E-06 | 8.25E-03 |
| muscle contraction (GO:0006936) | 248 | 10 | 1.61 | 6.23 | 6.83E-06 | 1.20E-02 |
| muscle system process (GO:0003012) | 298 | 12 | 1.93 | 6.22 | 8.05E-07 | 6.36E-03 |
| striated muscle cell differentiation (GO:0051146) | 199 | 8 | 1.29 | 6.21 | 5.97E-05 | 2.55E-02 |
| muscle cell differentiation (GO:0042692) | 240 | 9 | 1.55 | 5.79 | 3.46E-05 | 2.28E-02 |
| heart morphogenesis (GO:0003007) | 244 | 9 | 1.58 | 5.69 | 3.91E-05 | 2.21E-02 |
| striated muscle tissue development (GO:0014706) | 288 | 10 | 1.87 | 5.36 | 2.38E-05 | 1.98E-02 |
| muscle tissue development (GO:0060537) | 300 | 10 | 1.94 | 5.15 | 3.33E-05 | 2.39E-02 |
| muscle structure development (GO:0061061) | 467 | 14 | 3.02 | 4.63 | 2.80E-06 | 7.38E-03 |
| blood vessel morphogenesis (GO:0048514) | 404 | 11 | 2.62 | 4.2 | 7.95E-05 | 3.06E-02 |
| blood vessel development (GO:0001568) | 486 | 13 | 3.15 | 4.13 | 2.09E-05 | 1.84E-02 |
| vasculature development (GO:0001944) | 509 | 13 | 3.3 | 3.94 | 3.34E-05 | 2.30E-02 |
| cardiovascular system development (GO:0072358) | 519 | 13 | 3.36 | 3.87 | 4.06E-05 | 2.07E-02 |
| regulation of system process (GO:0044057) | 584 | 13 | 3.78 | 3.44 | 1.30E-04 | 4.45E-02 |
| negative regulation of transcription by RNA polymerase II (GO:0000122) | 836 | 18 | 5.42 | 3.32 | 9.52E-06 | 1.37E-02 |
| circulatory system development (GO:0072359) | 852 | 17 | 5.52 | 3.08 | 4.40E-05 | 2.05E-02 |
| animal organ morphogenesis (GO:0009887) | 930 | 18 | 6.02 | 2.99 | 3.81E-05 | 2.32E-02 |
| negative regulation of developmental process (GO:0051093) | 933 | 18 | 6.04 | 2.98 | 3.97E-05 | 2.17E-02 |
| cell-cell signaling (GO:0007267) | 1093 | 20 | 7.08 | 2.82 | 2.98E-05 | 2.36E-02 |
| negative regulation of nucleic acid-templated transcription (GO:1903507) | 1218 | 20 | 7.89 | 2.54 | 1.29E-04 | 4.55E-02 |
| negative regulation of RNA biosynthetic process (GO:1902679) | 1220 | 20 | 7.9 | 2.53 | 1.32E-04 | 4.36E-02 |
| negative regulation of nucleobase-containing compound metabolic process (GO:0045934) | 1406 | 23 | 9.11 | 2.53 | 4.02E-05 | 2.12E-02 |
| anatomical structure morphogenesis (GO:0009653) | 2102 | 31 | 13.62 | 2.28 | 1.17E-05 | 1.42E-02 |
| system process (GO:0003008) | 1937 | 28 | 12.55 | 2.23 | 7.15E-05 | 2.83E-02 |
| regulation of multicellular organismal process<br>(GO:0051239) | 3091 | 42 | 20.02 | 2.1 | 2.27E-06 | 7.19E-03 |

## References

1000 Genomes Project Consortium, Durbin, R.M., Altshuler, D.L., Durbin, R.M., Abecasis, G.R., Bentley, D.R., Chakravarti, A., Clark, A.G., Collins, F.S., De La Vega, F.M., et al. (2010). A map of human genome variation from population-scale sequencing. Nature 467, 1061–1073.

1000 Genomes Project Consortium, Auton, A., Brooks, L.D., Durbin, R.M., Garrison, E.P., Kang, H.M., Korbel, J.O., Marchini, J.L., McCarthy, S., McVean, G.A., et al. (2015). A global reference for human genetic variation. Nature 526, 68–74.

Allahyar, A., Vermeulen, C., Bouwman, B.A.M., Krijger, P.H.L., Verstegen, M.J.A.M., Geeven, G., van Kranenburg, M., Pieterse, M., Straver, R., Haarhuis, J.H.I., et al. (2018). Enhancer hubs and loop collisions identified from single-allele topologies. Nat. Genet. 50, 1151–1160.

Aouizerat, B.E., Vittinghoff, E., Musone, S.L., Pawlikowska, L., Kwok, P.-Y., Olgin, J.E., and Tseng, Z.H. (2011). GWAS for discovery and replication of genetic loci associated with sudden cardiac arrest in patients with coronary artery disease. BMC Cardiovasc. Disord. 11, 29.

Aragam, K.G., Chaffin, M., Levinson, R.T., McDermott, G., Choi, S.-H., Shoemaker, M.B., Haas, M.E., Weng, L.-C., Lindsay, M.E., Smith, J.G., et al. (2018). Phenotypic Refinement of Heart Failure in a National Biobank Facilitates Genetic Discovery. Circulation 139, 489–501.

Arking, D.E., Pulit, S.L., Crotti, L., van der Harst, P., Munroe, P.B., Koopmann, T.T., Sotoodehnia, N., Rossin, E.J., Morley, M., Wang, X., et al. (2014). Genetic association study of QT interval highlights role for calcium signaling pathways in myocardial repolarization. Nat. Genet. 46, 826–836.

Ashburner, M., Ball, C.A., Blake, J.A., Botstein, D., Butler, H., Cherry, J.M., Davis, A.P., Dolinski, K., Dwight, S.S., Eppig, J.T., et al. (2000). Gene ontology: tool for the unification of biology. The Gene Ontology Consortium. Nat. Genet. 25, 25–29.

Baxter, J.S., Leavy, O.C., Dryden, N.H., Maguire, S., Johnson, N., Fedele, V., Simigdala, N., Martin, L.A., Andrews, S., Wingett, S.W., et al. (2018). Capture Hi-C identifies putative target genes at 33 breast cancer risk loci. Nat. Commun.

Bhatia, S., and Kleinjan, D.A. (2014). Disruption of long-range gene regulation in human genetic disease: A kaleidoscope of general principles, diverse mechanisms and unique phenotypic consequences. Hum. Genet.

Bintu, B., Mateo, L.J., Su, J.-H., Sinnott-Armstrong, N.A., Parker, M., Kinrot, S., Yamaya, K., Boettiger, A.N., and Zhuang, X. (2018). Super-resolution chromatin tracing reveals domains and cooperative interactions in single cells. Science (80-.).

Bonev, B., Mendelson Cohen, N., Szabo, Q., Fritsch, L., Papadopoulos, G.L., Lubling, Y., Xu, X., Lv, X., Hugnot, J.-P., Tanay, A., et al. (2017). Multiscale 3D Genome Rewiring during Mouse Neural Development. Cell 171, 557–572.e24.

Buniello, A., MacArthur, J.A.L., Cerezo, M., Harris, L.W., Hayhurst, J., Malangone, C., McMahon, A., Morales, J., Mountjoy, E., Sollis, E., et al. (2019). The NHGRI-EBI GWAS Catalog of published genome-wide association studies, targeted arrays and summary statistics 2019. Nucleic Acids Res. 47, D1005– D1012.

Burren, O.S., Rubio García, A., Javierre, B.M., Rainbow, D.B., Cairns, J., Cooper, N.J., Lambourne, J.J., Schofield, E., Castro Dopico, X., Ferreira, R.C., et al. (2017). Chromosome contacts in activated T cells identify autoimmune disease candidate genes. Genome Biol.

Butler, A.M., Yin, X., Evans, D.S., Nalls, M.A., Smith, E.N., Tanaka, T., Li, G., Buxbaum, S.G., Whitsel, E.A., Alonso, A., et al. (2012). Novel loci associated with PR interval in a genome-wide association study of 10 African American cohorts. Circ. Cardiovasc. Genet. 5, 639–646.

Cairns, J., Freire-Pritchett, P., Wingett, S.W., Várnai, C., Dimond, A., Plagnol, V., Zerbino, D., Schoenfelder, S., Javierre, B.M., Osborne, C., et al. (2016). CHiCAGO: Robust detection of DNA looping interactions in Capture Hi-C data. Genome Biol.

Chang, C.C., Chow, C.C., Tellier, L.C., Vattikuti, S., Purcell, S.M., and Lee, J.J. (2015). Second-generation PLINK: rising to the challenge of larger and richer datasets. Gigascience 4, 7.

Chen, L., Jin, P., and Qin, Z.S. (2016). DIVAN: accurate identification of non-coding disease-specific risk variants using multi-omics profiles. Genome Biol. 17, 252.

Chesi, A., Wagley, Y., Johnson, M.E., Manduchi, E., Su, C., Lu, S., Leonard, M.E., Hodge, K.M., Pippin, J.A., Hankenson, K.D., et al. (2019). Genome-scale Capture C promoter interactions implicate effector genes at GWAS loci for bone mineral density. Nat. Commun. 10, 1260.

Cho, Y.S., Go, M.J., Kim, Y.J., Heo, J.Y., Oh, J.H., Ban, H.-J., Yoon, D., Lee, M.H., Kim, D.-J., Park, M.M.S., et al. (2009). A large-scale genome-wide association study of Asian populations uncovers genetic factors influencing eight quantitative traits. Nat. Genet. 41, 527–534.

Choy, M.K., Javierre, B.M., Williams, S.G., Baross, S.L., Liu, Y., Wingett, S.W., Akbarov, A., Wallace, C., Freire-Pritchett, P., Rugg-Gunn, P.J., et al. (2018). Promoter interactome of human embryonic stem cell-derived cardiomyocytes connects GWAS regions to cardiac gene networks. Nat. Commun.

Christophersen, I.E., Holmegard, H.N., Jabbari, J., Sajadieh, A., Haunsø, S., Tveit, A., Svendsen, J.H., and Olesen, M.S. (2013). Rare variants in GJA5 are associated with early-onset lone atrial fibrillation. Can. J. Cardiol. 29, 111–116.

Christophersen, I.E., Rienstra, M., Roselli, C., Yin, X., Geelhoed, B., Barnard, J., Lin, H., Arking, D.E., Smith, A. V, Albert, C.M., et al. (2017a). Large-scale analyses of common and rare variants identify 12 new loci associated with atrial fibrillation. Nat. Genet. 49, 946–952.

Christophersen, I.E., Magnani, J.W., Yin, X., Barnard, J., Weng, L.-C., Arking, D.E., Niemeijer, M.N., Lubitz, S.A., Avery, C.L., Duan, Q., et al. (2017b). Fifteen Genetic Loci Associated With the Electrocardiographic P Wave. Circ. Cardiovasc. Genet. 10.

Claussnitzer, M., Dankel, S.N., Kim, K.-H., Quon, G., Meuleman, W., Haugen, C., Glunk, V., Sousa, I.S., Beaudry, J.L., Puviindran, V., et al. (2015). FTO Obesity Variant Circuitry and Adipocyte Browning in Humans. N. Engl. J. Med. 373, 895–907.

Cornish, A.J., Hoang, P.H., Dobbins, S.E., Law, P.J., Chubb, D., Orlando, G., and Houlston, R.S. (2019). Identification of recurrent noncoding mutations in B-cell lymphoma using capture Hi-C. Blood Adv.

Dao, L.T.M., Galindo-Albarrán, A.O., Castro-Mondragon, J.A., Andrieu-Soler, C., Medina-Rivera, A., Souaid, C., Charbonnier, G., Griffon, A., Vanhille, L., Stephen, T., et al. (2017). Genome-wide characterization of mammalian promoters with distal enhancer functions. Nat. Genet. 49, 1073–1081.

Dekker, J., Rippe, K., Dekker, M., and Kleckner, N. (2002). Capturing Chromosome Conformation. Science (80-.). 295, 1306–1311.

Deng, W., Lee, J., Wang, H., Miller, J., Reik, A., Gregory, P.D., Dean, A., and Blobel, G.A. (2012). Controlling long-range genomic interactions at a native locus by targeted tethering of a looping factor. Cell 149, 1233–1244.

Dixon, J.R., Selvaraj, S., Yue, F., Kim, A., Li, Y., Shen, Y., Hu, M., Liu, J.S., and Ren, B. (2012). Topological domains in mammalian genomes identified by analysis of chromatin interactions. Nature 485, 376–380.

Dostie, J., Richmond, T.A., Arnaout, R.A., Selzer, R.R., Lee, W.L., Honan, T.A., Rubio, E.D., Krumm, A., Lamb, J., Nusbaum, C., et al. (2006). Chromosome Conformation Capture Carbon Copy (5C): a massively parallel solution for mapping interactions between genomic elements. Genome Res. 16, 1299–1309.

Durand, N.C., Shamim, M.S., Machol, I., Rao, S.S.P., Huntley, M.H., Lander, E.S., and Aiden, E.L. (2016). Juicer Provides a One-Click System for Analyzing Loop-Resolution Hi-C Experiments. Cell Syst. 3, 95–98.

Engreitz, J.M., Haines, J.E., Perez, E.M., Munson, G., Chen, J., Kane, M., McDonel, P.E., Guttman, M., and Lander, E.S. (2016). Local regulation of gene expression by lncRNA promoters, transcription and splicing. Nature.

Ernst, J., and Kellis, M. (2012). ChromHMM: automating chromatin-state discovery and characterization. Nat. Methods 9, 215–216.

Finn, E.H., Pegoraro, G., Brandão, H.B., Valton, A.L., Oomen, M.E., Dekker, J., Mirny, L., and Misteli, T. (2019). Extensive Heterogeneity and Intrinsic Variation in Spatial Genome Organization. Cell.

Gallagher, M.D., and Chen-Plotkin, A.S. (2018). The Post-GWAS Era: From Association to Function. Am. J. Hum. Genet. 102, 717–730.

Gasperini, M., Hill, A.J., McFaline-Figueroa, J.L., Martin, B., Kim, S., Zhang, M.D., Jackson, D., Leith, A., Schreiber, J., Noble, W.S., et al. (2019). A Genome-wide Framework for Mapping Gene Regulation via Cellular Genetic Screens. Cell.

Geeven, G., Teunissen, H., de Laat, W., and de Wit, E. (2018). peakC: a flexible, non-parametric peak calling package for 4C and Capture-C data. Nucleic Acids Res.

Gentleman, R.C., Carey, V.J., Bates, D.M., Bolstad, B., Dettling, M., Dudoit, S., Ellis, B., Gautier, L., Ge, Y., Gentry, J., et al. (2004). Bioconductor: open software development for computational biology and bioinformatics. Genome Biol. 5, R80.

Gollob, M.H., Jones, D.L., Krahn, A.D., Danis, L., Gong, X.-Q., Shao, Q., Liu, X., Veinot, J.P., Tang, A.S.L., Stewart, A.F.R., et al. (2006). Somatic mutations in the connexin 40 gene (GJA5) in atrial fibrillation. N. Engl. J. Med. 354, 2677–2688.

Gröschel, S., Sanders, M.A., Hoogenboezem, R., de Wit, E., Bouwman, B.A.M., Erpelinck, C., van der Velden, V.H.J., Havermans, M., Avellino, R., van Lom, K., et al. (2014). A single oncogenic enhancer rearrangement causes concomitant EVI1 and GATA2 deregulation in leukemia. Cell 157, 369–381.

GTEx Consortium, Gte. (2015). Human genomics. The Genotype-Tissue Expression (GTEx) pilot analysis: multitissue gene regulation in humans. Science 348, 648–660.

GTEx Consortium, J., Thomas, J., Salvatore, M., Phillips, R., Lo, E., Shad, S., Hasz, R., Walters, G., Garcia, F., Young, N., et al. (2013). The Genotype-Tissue Expression (GTEx) project. Nat. Genet. 45, 580–585.

Haarhuis, J.H.I., van der Weide, R.H., Blomen, V.A., Yáñez-Cuna, J.O., Amendola, M., van Ruiten, M.S., Krijger, P.H.L., Teunissen, H., Medema, R.H., van Steensel, B., et al. (2017). The Cohesin Release Factor WAPL Restricts Chromatin Loop Extension. Cell 169, 693–707.e14.

Harmston, N., Ing-Simmons, E., Perry, M., Barešić, A., and Lenhard, B. (2015). GenomicInteractions: An R/Bioconductor package for manipulating and investigating chromatin interaction data. BMC Genomics 16, 963.

van der Harst, P., and Verweij, N. (2018). Identification of 64 Novel Genetic Loci Provides an Expanded View on the Genetic Architecture of Coronary Artery Disease. Circ. Res. 122, 433–443.

den Hoed, M., Eijgelsheim, M., Esko, T., Brundel, B.J.J.M., Peal, D.S., Evans, D.M., Nolte, I.M., Segrè, A. V, Holm, H., Handsaker, R.E., et al. (2013). Identification of heart rate-associated loci and their effects on cardiac conduction and rhythm disorders. Nat. Genet. 45, 621–631.

Hughes, J.R., Roberts, N., McGowan, S., Hay, D., Giannoulatou, E., Lynch, M., De Gobbi, M., Taylor, S., Gibbons, R., and Higgs, D.R. (2014). Analysis of hundreds of cis-regulatory landscapes at high resolution in a single, high-throughput experiment. Nat. Genet. 46, 205–212.

Jäger, R., Migliorini, G., Henrion, M., Kandaswamy, R., Speedy, H.E., Heindl, A., Whiffin, N., Carnicer, M.J., Broome, L., Dryden, N., et al. (2015). Capture Hi-C identifies the chromatin interactome of colorectal cancer risk loci. Nat. Commun.

Javierre, B.M., Sewitz, S., Cairns, J., Wingett, S.W., Várnai, C., Thiecke, M.J., Freire-Pritchett, P., Spivakov, M., Fraser, P., Burren, O.S., et al. (2016). Lineage-Specific Genome Architecture Links Enhancers and Non-coding Disease Variants to Target Gene Promoters. Cell.

Kamburov, A., Stelzl, U., Lehrach, H., and Herwig, R. (2013). Nucleic Acids Research, 41, 793–800.

Kichaev, G., Bhatia, G., Loh, P.-R., Gazal, S., Burch, K., Freund, M.K., Schoech, A., Pasaniuc, B., and Price, A.L. (2019). Leveraging Polygenic Functional Enrichment to Improve GWAS Power. Am. J. Hum. Genet. 104, 65–75.

Kim, T.-K.K., and Shiekhattar, R. (2016). Diverse regulatory interactions of long noncoding RNAs. Curr. Opin. Genet. Dev. 36, 73–82.

Kishore, K., de Pretis, S., Lister, R., Morelli, M.J., Bianchi, V., Amati, B., Ecker, J.R., and Pelizzola, M. (2015). methylPipe and compEpiTools: a suite of R packages for the integrative analysis of epigenomics data. BMC Bioinformatics 16, 313.

Krijger, P.H.L., and de Laat, W. (2016). Regulation of disease-associated gene expression in the 3D genome. Nat. Rev. Mol. Cell Biol. 17, 771–782.

Krijger, P.H.L., and De Laat, W. (2013). Identical cells with different 3D genomes; cause and consequences? Curr. Opin. Genet. Dev.

Lambert, J.-C.C., Ibrahim-Verbaas, C.A., Harold, D., Naj, A.C., Sims, R., Bellenguez, C., DeStafano, A.L., Bis, J.C., Beecham, G.W., Grenier-Boley, B., et al. (2013). Meta-analysis of 74,046 individuals identifies 11 new susceptibility loci for Alzheimer’s disease. Nat. Genet. 45, 1452–1458.

de Lange, K.M., Moutsianas, L., Lee, J.C., Lamb, C.A., Luo, Y., Kennedy, N.A., Jostins, L., Rice, D.L., Gutierrez-Achury, J., Ji, S.-G., et al. (2017). Genome-wide association study implicates immune activation of multiple integrin genes in inflammatory bowel disease. Nat. Genet. 49, 256–261.

Li, G., Ruan, X., Auerbach, R.K., Sandhu, K.S., Zheng, M., Wang, P., Poh, H.M., Goh, Y., Lim, J., Zhang, J., et al. (2012a). Extensive Promoter-Centered Chromatin Interactions Provide a Topological Basis for Transcription Regulation. Cell 148, 84–98.

Li, G., Ruan, X., Auerbach, R.K., Sandhu, K.S., Zheng, M., Wang, P., Poh, H.M., Goh, Y., Lim, J., Zhang, J., et al. (2012b). Extensive promoter-centered chromatin interactions provide a topological basis for transcription regulation. Cell 148, 84–98.

Lieberman-Aiden, E., van Berkum, N.L., Williams, L., Imakaev, M., Ragoczy, T., Telling, A., Amit, I., Lajoie, B.R., Sabo, P.J., Dorschner, M.O., et al. (2009). Comprehensive mapping of long-range interactions reveals folding principles of the human genome. Science 326, 289–293.

Liu, N.Q., Ter Huurne, M., Nguyen, L.N., Peng, T., Wang, S.-Y., Studd, J.B., Joshi, O., Ongen, H., Bramsen, J.B., Yan, J., et al. (2017). The non-coding variant rs1800734 enhances DCLK3 expression through long-range interaction and promotes colorectal cancer progression. Nat. Commun. 8, 14418.

Locke, A.E., Kahali, B., Berndt, S.I., Justice, A.E., Pers, T.H., Day, F.R., Powell, C., Vedantam, S., Buchkovich, M.L., Yang, J., et al. (2015). Genetic studies of body mass index yield new insights for obesity biology. Nature 518, 197–206.

Love, M.I., Huber, W., and Anders, S. (2014). Moderated estimation of fold change and dispersion for RNA-seq data with DESeq2. Genome Biol. 15, 550.

Lupiáñez, D.G., Kraft, K., Heinrich, V., Krawitz, P., Brancati, F., Klopocki, E., Horn, D., Kayserili, H., Opitz, J.M., Laxova, R., et al. (2015). Disruptions of topological chromatin domains cause pathogenic rewiring of gene-enhancer interactions. Cell 161, 1012–1025.

Ma, J.-F., Yang, F., Mahida, S.N., Zhao, L., Chen, X., Zhang, M.L., Sun, Z., Yao, Y., Zhang, Y.-X., Zheng, G.-Y., et al. (2016). TBX5 mutations contribute to early-onset atrial fibrillation in Chinese and Caucasians. Cardiovasc. Res. 109, 442–450.

Marroni, F., Pfeufer, A., Aulchenko, Y.S., Franklin, C.S., Isaacs, A., Pichler, I., Wild, S.H., Oostra, B.A., Wright, A.F., Campbell, H., et al. (2009). A genome-wide association scan of RR and QT interval duration in 3 European genetically isolated populations: the EUROSPAN project. Circ. Cardiovasc. Genet. 2, 322– 328.

Martin, P., McGovern, A., Orozco, G., Duffus, K., Yarwood, A., Schoenfelder, S., Cooper, N.J., Barton, A., Wallace, C., Fraser, P., et al. (2015). Capture Hi-C reveals novel candidate genes and complex long-range interactions with related autoimmune risk loci. Nat. Commun.

Maurano, M.T., Humbert, R., Rynes, E., Thurman, R.E., Haugen, E., Wang, H., Reynolds, A.P., Sandstrom, R., Qu, H., Brody, J., et al. (2012). Systematic localization of common disease-associated variation in regulatory DNA. Science (80-.). 337, 1190–1195.

Mifsud, B., Tavares-Cadete, F., Young, A.N., Sugar, R., Schoenfelder, S., Ferreira, L., Wingett, S.W., Andrews, S., Grey, W., Ewels, P.A., et al. (2015). Mapping long-range promoter contacts in human cells with high-resolution capture Hi-C. Nat. Genet. 47, 598–606.

Montefiori, L.E., Sobreira, D.R., Sakabe, N.J., Aneas, I., Joslin, A.C., Hansen, G.T., Bozek, G., Moskowitz, I.P., McNally, E.M., and Nóbrega, M.A. (2018). A promoter interaction map for cardiovascular disease genetics. Elife.

Mumbach, M.R., Satpathy, A.T., Boyle, E.A., Dai, C., Gowen, B.G., Cho, S.W., Nguyen, M.L., Rubin, A.J., Granja, J.M., Kazane, K.R., et al. (2017). Enhancer connectome in primary human cells identifies target genes of disease-associated DNA elements. Nat. Genet. 49, 1602–1612.

Nadadur, R.D., Broman, M.T., Boukens, B., Mazurek, S.R., Yang, X., van den Boogaard, M., Bekeny, J., Gadek, M., Ward, T., Zhang, M., et al. (2016). Pitx2 modulates a Tbx5-dependent gene regulatory network to maintain atrial rhythm. Sci. Transl. Med. 8, 354ra115.

Nora, E.P., Lajoie, B.R., Schulz, E.G., Giorgetti, L., Okamoto, I., Servant, N., Piolot, T., van Berkum, N.L., Meisig, J., Sedat, J., et al. (2012). Spatial partitioning of the regulatory landscape of the X-inactivation centre. Nature 485, 381–385.

Nuebler, J., Fudenberg, G., Imakaev, M., Abdennur, N., and Mirny, L.A. (2018). Chromatin organization by an interplay of loop extrusion and compartmental segregation. Proc. Natl. Acad. Sci. 115, E6697– E6706.

Olivares-Chauvet, P., Mukamel, Z., Lifshitz, A., Schwartzman, O., Elkayam, N.O., Lubling, Y., Deikus, G., Sebra, R.P., and Tanay, A. (2016). Capturing pairwise and multi-way chromosomal conformations using chromosomal walks. Nature, 540, 296–300.

Orlando, G., Law, P.J., Cornish, A.J., Dobbins, S.E., Chubb, D., Broderick, P., Litchfield, K., Hariri, F., Pastinen, T., Osborne, C.S., et al. (2018). Promoter capture Hi-C-based identification of recurrent noncoding mutations in colorectal cancer. Nat. Genet.

Prins, B.P., Mead, T.J., Brody, J.A., Sveinbjornsson, G., Ntalla, I., Bihlmeyer, N.A., van den Berg, M., Bork-Jensen, J., Cappellani, S., Van Duijvenboden, S., et al. (2018). Exome-chip meta-analysis identifies novel loci associated with cardiac conduction, including ADAMTS6. Genome Biol. 19, 87.

R Core Team (2015). R: A Language and Environment for Statistical Computing.

Rao, S.S.P., Huntley, M.H., Durand, N.C., Stamenova, E.K., Bochkov, I.D., Robinson, J.T., Sanborn, A.L., Machol, I., Omer, A.D., Lander, E.S., et al. (2014). A 3D Map of the Human Genome at Kilobase Resolution Reveals Principles of Chromatin Looping. Cell 159, 1665–1680.

Rao, S.S.P., Huang, S.-C., Glenn St Hilaire, B., Engreitz, J.M., Perez, E.M., Kieffer-Kwon, K.-R., Sanborn, A.L., Johnstone, S.E., Bascom, G.D., Bochkov, I.D., et al. (2017). Cohesin Loss Eliminates All Loop Domains. Cell 171, 305–320.e24.

Roadmap Epigenomics Consortium, A., Kundaje, A., Meuleman, W., Ernst, J., Bilenky, M., Yen, A., Heravi-Moussavi, A., Kheradpour, P., Zhang, Z., Wang, J., et al. (2015). Integrative analysis of 111 reference human epigenomes. Nature 518, 317–330.

Robinson, J.T., Turner, D., Durand, N.C., Thorvaldsdóttir, H., Mesirov, J.P., and Aiden, E.L. (2018). Juicebox.js Provides a Cloud-Based Visualization System for Hi-C Data. Cell Syst. 6, 256–258.e1.

Roselli, C., Chaffin, M.D., Weng, L.-C., Aeschbacher, S., Ahlberg, G., Albert, C.M., Almgren, P., Alonso, A., Anderson, C.D., Aragam, K.G., et al. (2018). Multi-ethnic genome-wide association study for atrial fibrillation. Nat. Genet. 50, 1225–1233.

Ruf, S., Symmons, O., Uslu, V.V., Dolle, D., Hot, C., Ettwiller, L., and Spitz, F. (2011). Large-scale analysis of the regulatory architecture of the mouse genome with a transposon-associated sensor. Nat. Genet. 43, 379–386.

Schaub, M.A., Boyle, A.P., Kundaje, A., Batzoglou, S., and Snyder, M. (2012). Linking disease associations with regulatory information in the human genome. Genome Res. 22, 1748–1759.

Schizophrenia Working Group of the Psychiatric Genomics Consortium (2014). Biological insights from 108 schizophrenia-associated genetic loci. Nature 511, 421–427.

Schmidt, E.M., Zhang, J., Zhou, W., Chen, J., Mohlke, K.L., Chen, Y.E., and Willer, C.J. (2015). GREGOR: evaluating global enrichment of trait-associated variants in epigenomic features using a systematic, data-driven approach. Bioinformatics 31, 2601–2606.

Schwarzer, W., Abdennur, N., Goloborodko, A., Pekowska, A., Fudenberg, G., Loe-Mie, Y., Fonseca, N.A., Huber, W., H Haering, C., Mirny, L., et al. (2017). Two independent modes of chromatin organization revealed by cohesin removal. Nature 551, 51–56.

Scott, R.A., Scott, L.J., Mägi, R., Marullo, L., Gaulton, K.J., Kaakinen, M., Pervjakova, N., Pers, T.H., Johnson, A.D., Eicher, J.D., et al. (2017). An Expanded Genome-Wide Association Study of Type 2 Diabetes in Europeans. Diabetes 66, 2888–2902.

van Setten, J., Brody, J.A., Jamshidi, Y., Swenson, B.R., Butler, A.M., Campbell, H., Del Greco, F.M., Evans, D.S., Gibson, Q., Gudbjartsson, D.F., et al. (2018). PR interval genome-wide association meta-analysis identifies 50 loci associated with atrial and atrioventricular electrical activity. Nat. Commun. 9, 2904.

Sexton, T., Yaffe, E., Kenigsberg, E., Bantignies, F., Leblanc, B., Hoichman, M., Parrinello, H., Tanay, A., and Cavalli, G. (2012). Three-dimensional folding and functional organization principles of the Drosophila genome. Cell 148, 458–472.

Simonis, M., Klous, P., Splinter, E., Moshkin, Y., Willemsen, R., de Wit, E., van Steensel, B., and de Laat, W. (2006). Nuclear organization of active and inactive chromatin domains uncovered by chromosome conformation capture-on-chip (4C). Nat. Genet. 38, 1348–1354.

Smemo, S., Tena, J.J., Kim, K.-H., Gamazon, E.R., Sakabe, N.J., Gómez-Marín, C., Aneas, I., Credidio, F.L., Sobreira, D.R., Wasserman, N.F., et al. (2014). Obesity-associated variants within FTO form long-range functional connections with IRX3. Nature 507, 371–375.

Speliotes, E.K., Willer, C.J., Berndt, S.I., Monda, K.L., Thorleifsson, G., Jackson, A.U., Lango Allen, H., Lindgren, C.M., Luan, J., Mägi, R., et al. (2010). Association analyses of 249,796 individuals reveal 18 new loci associated with body mass index. Nat. Genet. 42, 937–948.

Spielmann, M., Lupiáñez, D.G., and Mundlos, S. (2018). Structural variation in the 3D genome. Nat. Rev. Genet. 19, 453–467.

The Gene Ontology Consortium (2017). Expansion of the Gene Ontology knowledgebase and resources. Nucleic Acids Res. 45, D331–D338.

Thomas, P.D., Kejariwal, A., Guo, N., Mi, H., Campbell, M.J., Muruganujan, A., and Lazareva-Ulitsky, B. (2006). Applications for protein sequence-function evolution data: mRNA/protein expression analysis and coding SNP scoring tools. Nucleic Acids Res. 34, W645–W650.

Tolhuis, B., Palstra, R.J., Splinter, E., Grosveld, F., and de Laat, W. (2002). Looping and interaction between hypersensitive sites in the active beta-globin locus. Mol. Cell 10, 1453–1465.

Turner, S.D. (2018). qqman: an R package for visualizing GWAS results using Q-Q and manhattan plots Software • Review • Repository • Archive.

Verweij, N., Mateo Leach, I., van den Boogaard, M., van Veldhuisen, D.J., Christoffels, V.M., LifeLines Cohort Study, H.L., Hillege, H.L., van Gilst, W.H., Barnett, P., de Boer, R.A., et al. (2014). Genetic determinants of P wave duration and PR segment. Circ. Cardiovasc. Genet. 7, 475–481.

Wang, Y., Song, F., Zhang, B., Zhang, L., Xu, J., Kuang, D., Li, D., Choudhary, M.N.K., Li, Y., Hu, M., et al. (2018). The 3D Genome Browser: a web-based browser for visualizing 3D genome organization and long-range chromatin interactions. Genome Biol. 19, 151.

Warren, H.R., Evangelou, E., Cabrera, C.P., Gao, H., Ren, M., Mifsud, B., Ntalla, I., Surendran, P., Liu, C., Cook, J.P., et al. (2017). Genome-wide association analysis identifies novel blood pressure loci and offers biological insights into cardiovascular risk. Nat. Genet. 49, 403–415.

Van De Werken, H.J.G., De Vree, P.J.P., Splinter, E., Holwerda, S.J.B., Klous, P., De Wit, E., and De Laat, W. (2012). 4C technology: Protocols and data analysis. Methods Enzymol.

Wild, P.S., Felix, J.F., Schillert, A., Teumer, A., Chen, M.-H., Leening, M.J.G.G., Völker, U., Großmann, V., Brody, J.A., Irvin, M.R., et al. (2017). Large-scale genome-wide analysis identifies genetic variants associated with cardiac structure and function. J. Clin. Invest. 127, 1798–1812.

Willer, C.J., Schmidt, E.M., Sengupta, S., Peloso, G.M., Gustafsson, S., Kanoni, S., Ganna, A., Chen, J., Buchkovich, M.L., Mora, S., et al. (2013). Discovery and refinement of loci associated with lipid levels. Nat. Genet. 45, 1274–1283.

Wingett, S., Ewels, P., Furlan-Magaril, M., Nagano, T., Schoenfelder, S., Fraser, P., and Andrews, S. (2015). HiCUP: pipeline for mapping and processing Hi-C data. F1000Research, 4, 1310.

Wray, N.R., Ripke, S., Mattheisen, M., Trzaskowski, M., Byrne, E.M., Abdellaoui, A., Adams, M.J., Agerbo, E., Air, T.M., Andlauer, T.M.F., et al. (2018). Genome-wide association analyses identify 44 risk variants and refine the genetic architecture of major depression. Nat. Genet. 50, 668–681.

Zhang, Y., Liu, T., Meyer, C.A., Eeckhoute, J., Johnson, D.S., Bernstein, B.E., Nusbaum, C., Myers, R.M., Brown, M., Li, W., et al. (2008). Model-based analysis of ChIP-Seq (MACS). Genome Biol. 9, R137.

Zhang, Y., McCord, R.P., Ho, Y.-J., Lajoie, B.R., Hildebrand, D.G., Simon, A.C., Becker, M.S., Alt, F.W., and Dekker, J. (2012). Spatial organization of the mouse genome and its role in recurrent chromosomal translocations. Cell 148, 908–921.

Ziebarth, J.D., Bhattacharya, A., and Cui, Y. (2013). CTCFBSDB 2.0: A database for CTCF-binding sites and genome organization. Nucleic Acids Res.

